# Visual category representations in the infant brain

**DOI:** 10.1101/2021.11.03.466293

**Authors:** Siying Xie, Stefanie Hoehl, Merle Moeskops, Ezgi Kayhan, Christian Kliesch, Bert Turtleton, Moritz Köster, Radoslaw M. Cichy

## Abstract

Visual categorization is a human core cognitive capacity ^1,2^ that depends on the development of visual category representations in the infant brain ^3–7^. However, the exact nature of infant visual category representations and their relationship to the corresponding adult form remains unknown ^8^. Our results clarify the nature of visual category representations from electroencephalography (EEG) data in 6- to 8-month-old infants and their developmental trajectory towards adult maturity in the key characteristics of temporal dynamics ^2,9^, representational format ^10–12^, and spectral properties ^13,14^. Temporal dynamics change from slowly emerging, developing representations in infants to quickly emerging, complex representations in adults. Despite those differences, infants and adults already partly share visual category representations. The format of infants’ representations is visual features of low to intermediate complexity, whereas adults’ representations also encode high complexity features. Theta band activity contributes to visual category representations in infants, and these representations are shifted to the alpha/beta band in adults. Together, we reveal the developmental neural basis of visual categorization in humans, show how information transmission channels change in development, and demonstrate the power of advanced multivariate analysis techniques in infant EEG research for theory building in developmental cognitive science.

## RESULTS & DISCUSSION

The ability to recognize and categorize visual objects effortlessly and within the blink of an eye is a core human cognitive capacity ^1,2^ that develops through learning and interaction with the environment. Behavioral research in infants using looking times ^6,7,15^ and neural markers of attention provides evidence for visual category processing ^16^ and learning ^5,17^ already within the first year of life.

In adults fundamental research in human and non-human primates has described the nature of the neural representations underlying mature visual categorization abilities, revealing their temporal dynamics ^2,9^, what features they encode ^10–12^, their cortical locus ^11,18^, and how they relate to neural oscillations ^13,14^. In contrast, these key characteristics of visual category representations ^19–23^ are less well understood in infants due to strong methodological challenges in human and non-human infant neuroimaging research ^8,24,25^. In particular, research using EEG – the workhorse of infant neuroimaging for decades – has yielded insights that are principally limited in two ways. One research approach focused on assessing the successful outcome of visual categorization rather than the underlying representations themselves ^26^. Thus, the insights gained about representations are indirect. Another research approach did assess underlying representations directly but was limited to the category of faces ^3,27^ for which known neural markers exist. Thus, the generalizability from the unique and small stimulus subset to the broad set of visual categories of the visual world remains unclear.

Here we overcome this double impasse to reveal the nature of general visual category representations for various object categories in 6- to 8-month-old infants from EEG data. We do so by leveraging an integrated multivariate analysis framework of multivariate classification ^9^ and direct quantitative comparison ^28^ of the infant to adult EEG data and deep learning models of vision ^12^.

### The temporal dynamics of visual category representations

Infant participants (*n* = 40) viewed 128 images of real-world objects from four categories (i.e., toys, bodies, houses and faces, see **Figure 1A;** for rationale of category choice see **Method Details**) while we acquired EEG data. The age group was chosen based on extensive work showing that by this age infants discriminate between basic level categories reliably ^26,29–31^. Images were presented for 2s every 2.7–2.9s. For direct comparison we acquired EEG data in adult participants (*n* = 20) viewing the same stimulus set with an adapted experimental design (**Figure S1A,B**). We consider the epoch of −100ms to +1,000ms with respect to stimulus onset in our analyses.

**Figure 1.**
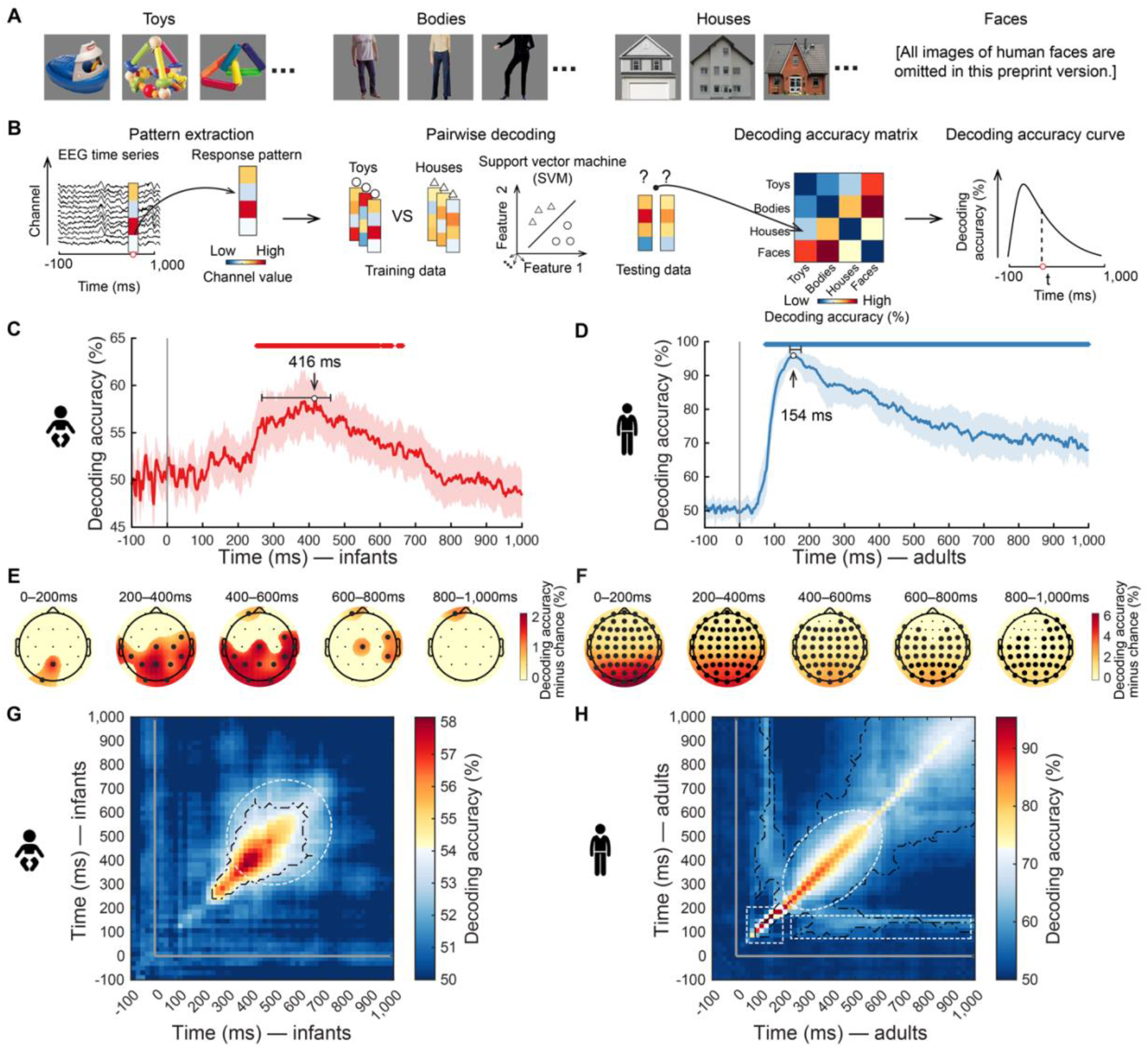
Experimental design and results of time-resolved multivariate analysis. **(A)** The stimulus set comprised 32 cut-out images from four categories each: toys, headless bodies, houses, and faces (full set see **OSF Repository**). **(B)** Time-resolved multivariate analysis on EEG data. First, we extracted condition-specific EEG sensor activation values for every time point in the epoch and formed them into response vectors. Then, using a leave-one-out cross-validation scheme, we trained and tested a support vector machine to classify visual object categories from the response vectors. The results (pairwise decoding accuracy, 50% chance level) were aggregated in a decoding accuracy matrix of size 4 × 4, indexed in rows and columns by the conditions classified. The matrix is symmetric along the diagonal, and the diagonal is undefined. Averaging the lower triangular part of the matrix resulted in grand average decoding accuracy as a measure of how well visual representations discriminate categories at a particular time point. **(C,D)** The grand average time course of visual category decoding in infants **(C)** and adults **(D)**. The gray vertical line indicates onset of image presentation. Shaded margins indicate 95% confidence intervals (CIs) of decoding accuracy. Horizontal error bars indicate 95% CIs of peak latency. Rows of asterisks indicate time points with significantly above-chance decoding accuracy (infant *n* = 40 or adult *n* = 20, right-tailed sign permutation tests, cluster-defining threshold *P* < .005, corrected significance level *P* < .05). Detailed statistical information is listed in **Table S1A**. For visualization of single participant infant data see **GitHub Repository: Visualization**. **(E,F)** Results of category classification in EEG channel-space searchlight analysis for infants **(E)** and adults **(F)**. Bold dots indicate the EEG channels with significantly above-chance decoding accuracy (right-tailed sign-permutation tests, *P* < .05, FDR-corrected). **(G,H)** Results of time-generalization analysis for infants **(G)** and adults **(H)**. Detailed statistical information is listed in **Table S1E**. For visualization of single participant infant data see **GitHub Repository: Visualization**. The gray vertical and horizontal lines indicate the onsets of image presentation. Black outlines indicate time-point combinations with significantly above-chance decoding accuracy (right-tailed sign permutation tests, cluster-defining threshold *P* < .005, corrected significance level *P* < .05). See also **Figure S1** and **Table S1**.

To reveal the time course with which visual category is discriminated by visual representations we used time-resolved multivariate pattern analysis ^9^ (**Figure 1B**). We report peak latency (95% confidence intervals (CIs) in brackets) as the time point during neural processing when category information was most explicit, as well as onset and offset of significance for each group.

In infants (**Figure 1C**), the classification curve rose gradually from 100ms onwards, reaching significance at 252ms (250–254ms), followed by a broad peak at 416ms (268–462ms) and a gradual decline. This pattern of result did not depend on any particular object or category (**Figure S1E,F**), held equally for classifications within and across the animacy division **(Figure S1G,H**), and emerged equivalently for alternative common analysis schemes (**Figure S1I–K**). In contrast, in adults, the classification curve had a different shape (**Figure 1D**). It emerged earlier (significant at 72ms (72–74ms)) and faster, peaking at 154ms (144–176ms) than in infants (*P* < .001, bootstrap test, **Table S1A**). The observed delay is not only due to longer latencies already at the early cortical processing stages: the P1 component peak in infants was delayed by 22–68ms in infants (**Figure S1C,D**, **Table S1F**), consistent with previous studies ^6,32–34^. Instead, the grand average ERP peak was much stronger, delayed by 98–242ms. This suggests that the observed peak latency differences with which category representations emerge reflect a mixture of processing delays at early and late processing stages.

Searchlight analysis in EEG channel space revealed that information about visual category representations was highest in EEG channels overlying occipitoparietal cortex in both infants (**Figure 1E**) and adults (**Figure 1F**), tentatively suggesting partly similar cortical sources in the posterior cortex.

This multivariate approach constitutes a novel analytical access point to visual category representations in infants from EEG data. Noteworthy, there is no simple mapping function of our results to the results of classical univariate results, as the approaches differ in many aspects. Univariate analyses focus on single electrodes or averages, while multivariate analyses focus on patterns across electrodes, potentially increasing sensitivity ^35,36^. In adults, univariate and multivariate analyses also do not directly agree ^9,37^. However, the multivariate results carry meaning, as they can be meaningfully related to behavior ^9,38^. Further research combining multivariate analyses with behavioral measures ^7,15^ in infants is needed.

The rise and fall of the classification curves in a few hundred milliseconds might indicate rapid changes in the underlying visual representations ^2,9^, slow ramping up of persistent representations or a combination of both. To investigate this, we assessed the temporal stability of visual representations using time-generalization analysis ^39^. We determined how well classifiers trained on predicting visual category from EEG at one time point perform when tested at other time points. Lack of generalization across time indicates transience of the underlying visual representation, whereas generalization across time indicates persistence.

In both infants (**Figure 1G**) and adults (**Figure 1H**), classification accuracy was highest along the diagonal (i.e., similar time points for training and testing) with a broadening over time (white dotted ellipse). This result suggests common neural mechanisms of a rapid sequence of processing steps that result in an outcome held online for further use, indicated by rapidly changing transient representations at earlier time points and more slowly changing persistent representations at later time points, respectively.

In addition to this general similarity between infants and adults, two notable differences were indicative of incomplete development of feedforward and feedback information processing in infants. For one, early after stimulus onset, when neural processing is dominantly feedforward, in adults, we observed high classification accuracy trailing the diagonal narrowly (**Figure 1H**, 50–200ms, dotted square), indicating rapid changing representations. Infants did not exhibit such signals. This pattern suggests incomplete development of feedforward visual information processing mechanisms in infants. Secondly, in adults, the classifier generalized well for the time point combination of 100–200ms and 200–1,000ms (**Figure 1H**, white striped rectangle). This suggests highly persistent representations, likely emerging in the early visual cortex ^9^. There were no such signals in infants. This indicates incomplete neural structures for recurrent processing that maintain visual information online for long stretches of time.

While the overall pattern of results did not depend on any particular category (**Figure S1L,M**), we cannot exclude that differences between age groups could also be due to differences in experimental task or signal-to-noise ratio (SNR). We boost SNR in adults compared to infants by design to increase the chance of identifying similarities at the cost of interpretative difficulties for differences. These difficulties are, however, alleviated by focusing on peaks as core measures for interpretation whose size, but not latency depends on SNR.

### Shared visual category representations between infants and adults

The results so far show that we identified visual category representations in both infants and adults and that their time courses have both similar and different aspects. However, we have not tested whether infants and adults have similar category representations. An alternative hypothesis is that we observe time courses of category classification for infants and adults, but those are unrelated rather than shared representations. Direct identification of shared representations between infants and adults is challenging due to differences in the time course over which the representations emerge and the EEG channel spaces differ. We used a time-generalization variant of representational similarity analysis (RSA) ^28^ (**Figure 2A**) to overcome these hurdles. In short, we abstracted multivariate signals from the incommensurate infant and adult EEG channel spaces to a common representational dissimilarity space, and we compared the signals across all time point combinations.

**Figure 2.**
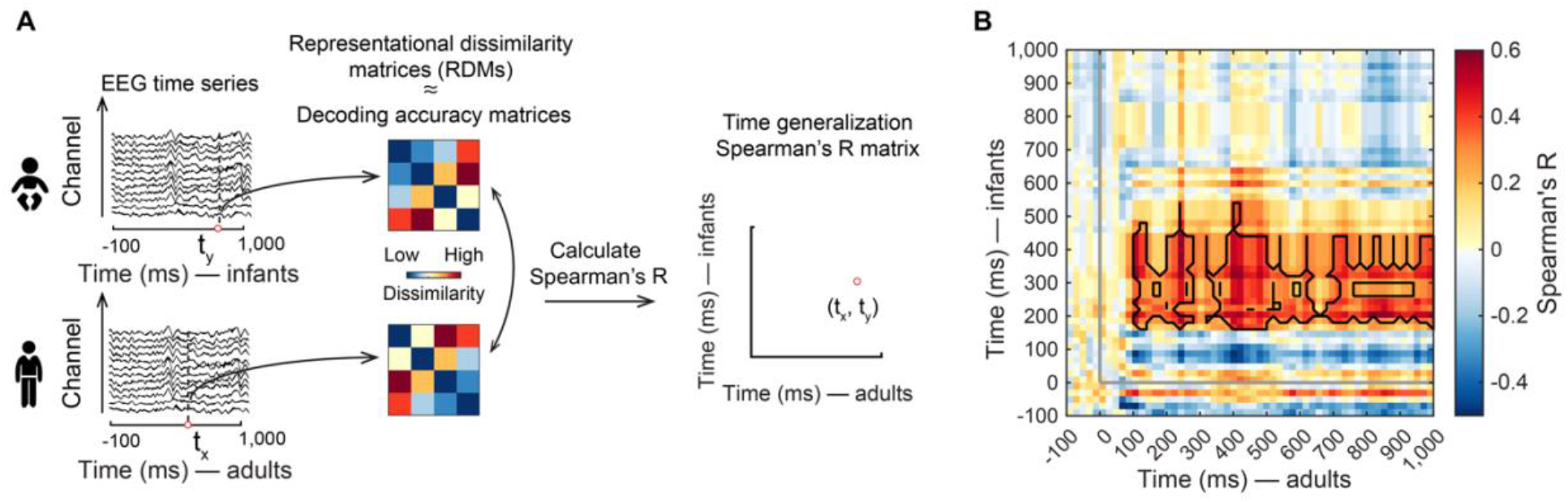
Category representations shared between infants and adults. **(A)** We used RSA to relate category representations in infants and adults. We interpret decoding accuracy as a dissimilarity measure on the assumption that the more dissimilar two representations are, the better the classifier performs. This allowed us to use time-resolved decoding accuracy matrices as representational dissimilarity matrices (RDMs) that summarize representational similarities between category representations. We compared RDMs (Spearman’s *R*) in infants (average across participants) and adults (for each participant separately) for all time point combinations (t_x_, t_y_), assigning the values to a time-generalization matrix indexed in rows and columns by the time in adults (t_x_) and infants (t_y_). **(B)** Average time-generalization matrix relating category representations in infants and adults over time. Detailed statistical information is listed in **Table S2A**. For visualization of single participant data see **GitHub Repository: Visualization**. The gray lines indicate image onset. Black outlines indicate time point combinations with significant correlation (*n* = 20, right-tailed sign permutation tests, cluster-defining threshold *P* < .005, corrected significance level *P* < .05). See also **Figure S2** and **Table S2**.

We observed similarity in visual category representations between infants and adults at the time point combinations of 160–540ms in infants, and 100–1,000ms in adults (**Figure 2B**, peak latency in infants: 200ms (200–360ms); in adults: 120ms (120–1,000ms)). This result was similarly achieved for alternative processing and data aggregation choices (**Figure S2A,B**), and did not depend on any particular category except on toys (**Figure S2C**). Our findings establish quantitatively and directly that infants and adults share visual category representations.

In sum, the emerging picture is one of not yet fully developed dynamics of adult-like visual category representations in infants. Representations in infants emerged later, slower, and lacked particular components of feedforward and recurrent processing, possibly related to immature myelination ^40^ and synaptic connectivity ^41^. Nevertheless, representations in infants and adults shared large-scale temporal dynamics that encoded visual category information similarly, consistent with previous showing partly adult-like behavioral ^3,5,7,15^ and neural ^19,20^ category sensitivity in the first year of age.

Our approach goes beyond previous EEG work in developmental visual neuroscience in three ways. First, rather than relying on indirect inference from attentional markers indicating successful categorization ^26^, our approach assessed representations directly as they emerge with millisecond resolution. Second, our approach is not limited to the face category and face-specific EEG components ^3,27^, but allows the study of potentially any visual category. Third, our approach enabled a new quantitative comparison ^28^ of infant and adult visual category representations.

Our results make direct predictions for the detailed developmental trajectory of visual category representations ^42^. We expect category representations to emerge increasingly earlier and with faster temporal dynamics with increasing age, with additional feedforward and feedback components appearing at critical stages until a mature adult-like system emerges. Our approach makes these predictions immediately testable in future studies using other age groups between early infancy and adulthood.

More broadly, our multivariate EEG analysis approach demonstrates a novel access point to largely unmapped neural representations in the infant brain, with strong potential to inform theories of cognitive development for cognitive capacities that emerge in the first year of life, such as object learning ^5,6,17^, speech processing ^43^, and core knowledge systems ^44^. Combined with human infant fMRI ^19,20,25^ and behavioral assessment ^15^ in a common framework ^45^, this promises to reveal the unknown spatiotemporal neural dynamics underlying cognitive functions in infants in the future.

### The format of visual category representations

The time-resolved multivariate pattern analysis revealed the presence and dynamics of visual category representations in the infant and adult brain. However, by itself, it is unable to specify their format, i.e., what type of visual features they encode. We hypothesized that adults would encode visual features represented at all levels of the visual processing hierarchy from low-to high complexity ^46^. Instead, infants would encode visual features rather of low- and mid-complexity, as predicted from visual behavior ^7,26^ and anatomical development patterns ^42^ of the infant visual brain.

To determine the format of category representations, we related them to computational models of vision (**Figure 3A**). We probed two types of models: a Gabor wavelet pyramid model as a model of simple visual features ^47^, and the deep neural network VGG-19 model ^48^ trained on object categorization, which exhibits a hierarchy of low-to-high complexity features along with its layers, and predicts activity along the visual processing hierarchy of the adult human brain well ^12,49^.

**Figure 3.**
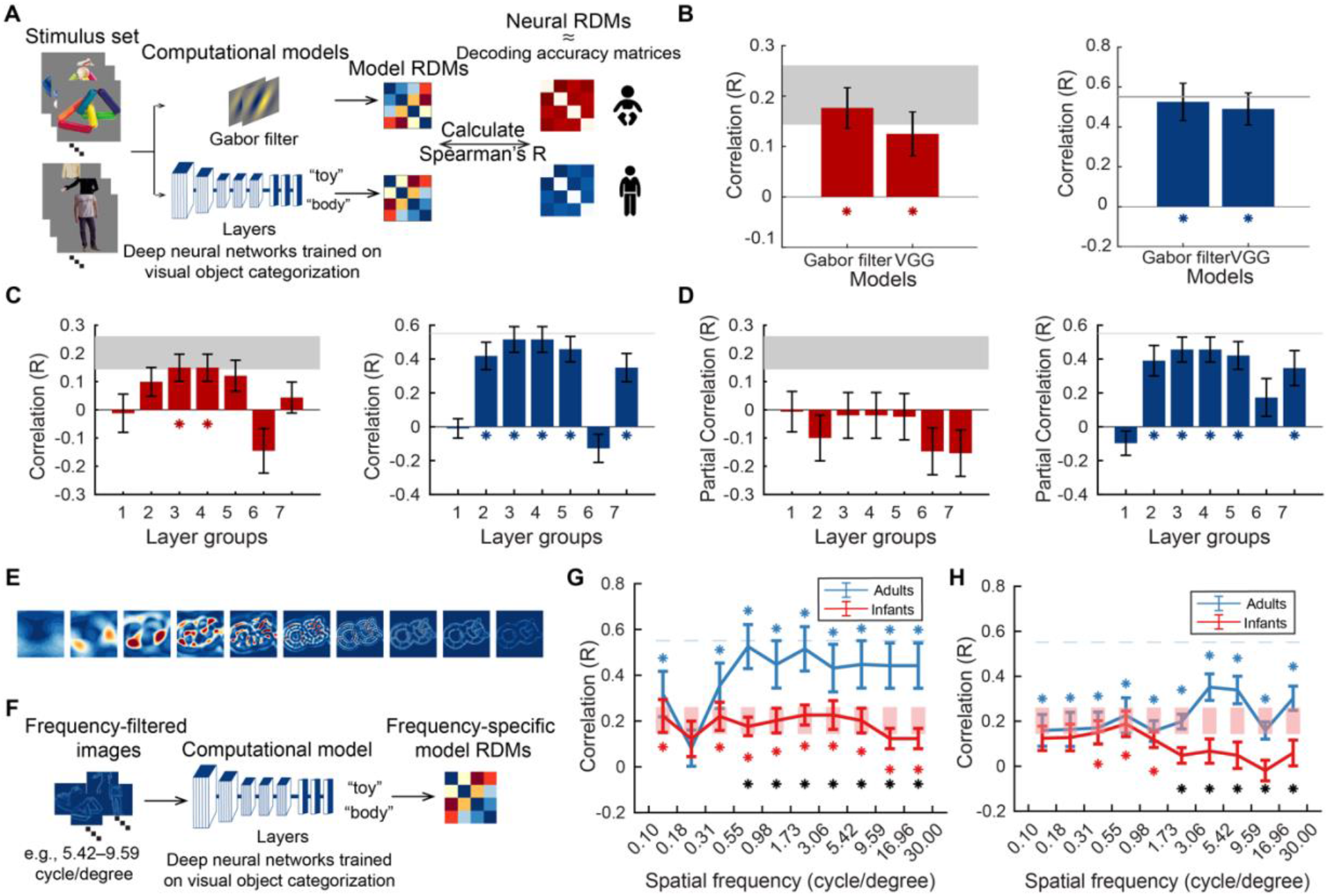
The format of category representations in infants and adults. **(A)** We characterized what type of visual features are encoded in category representations in infants and adults by relating them to computational models using RSA. We ran the stimulus images through a Gabor filter model and the VGG-19 deep neural network trained on object categorization. We constructed RDMs from their unit activation patterns (visualized in **GitHub Repository: Visualization**). We then compared model RDMs to infant and adult neural RDMs (constructed as the average of RDMs over time from 95% CIs around peak latency of time-resolved category classification, see **Figure 1C,D**). **(B)** Results for infants (left) and adults (right) at the whole-model level. Asterisks indicate significant correlation. **(C)** Results for infants (left) and adults (right) at the DNN layer level. For **(B,C)**, error bars represent standard errors of the mean. Asterisks indicate significant correlation (infant *n* = 40, adult *n* = 20, two-tailed sign-permutation tests, *P* < .05, FDR-corrected). Statistical details (i.e., correlations and *P*-values) are in **Table S3A**. **(D)** Results for infants (left) and adults (right) at the DNN layer level after removing the effect of the other age group respectively by partialling out the average RDM. Error bars represent standard errors of the mean. Asterisks indicate significant correlation. **(E)** Example of Butterworth filtered images in different spatial frequencies. **(F)** We characterized visual features encoded in visual representations in terms of spatial frequency content. We ran the frequency-filtered images through the VGG-19 DNN. We constructed spatial-frequency-specific RDMs from the DNN unit activation patterns. We then compared model RDMs to infant and adult neural RDMs as described in **(A)**. **(G)** Relating frequency-specific image content to neural representations. The results indicate a significant correlation across all frequencies (except one bin at 0.18 cycle per degree (cpd)) in both infants and adults, with higher correlations for adults than infants above 1 cpd. **(H)** Spatial-frequency-specific results for infants (blue curve) and adults (red curve) at the whole-model level. For **(G,H)**, asterisks color-coded as result curves indicate statistical significance (infant *n* = 40, adult *n* = 20, two-tailed sign-permutation tests, *P* < .05, FDR-corrected); black asterisks indicate significant difference between age groups (two-tailed Mann-Whitney U tests, *P* < .05, FDR-corrected). See also **Figure S3** and **Table S3**.

Assessing first the Gabor model and an aggregated summary of the VGG model across layers, we found similar representations between both models and infant and adult visual representations (**Figure 3B**). This suggested that features ranging from low to high complexity might contribute, and invited further in-depth analysis.

Turning to the VGG model first we conducted a finer investigation of VGG at the level of layers. Considering each layer separately, we found that in infants, middle layers predicted brain activity best, with layer groups 3 and 4 being significant (**Figure 3C**, left). In contrast, layers at all stages were significantly predictive in adults (**Figure 3C**, right). This pattern of results was also achieved for other types of deep neural network architectures (**Figure S3A,B**) and independent of data selection choices (**Figure S3C,D**), demonstrating the robustness of the result. This pattern of results suggests that infants and adults share similar visual features with the VGG model at intermediate complexity. We ascertained this in two ways. First, using partial correlation, we related the VGG model to each age group while partialling out the effect of the other age group. This abolished all effects in infants (**Figure 3D**, left) while leaving the resulting pattern in adults unchanged (**Figure 3D**, right), suggesting that the features underlying visual category representation in infants are a subset of the features in adults. Second, we conducted a variance partitioning analysis between the VGG model and infant and adult visual representations at layers 3 and 4, revealing shared variance (both R^2^ = 0.16; *P* < .05, FDR-corrected). This reveals that in infants category representations are in the format of low to intermediate complexity features and form a subset of the representations seen in adults, while in adults category is discriminated by features at all levels of complexity.

The prediction by the Gabor filter model and by the early layers of VGG in adults suggests that in both age groups, the category is discriminated by representations encoding features not only of low- and intermediate, but also low complexity, albeit to a different degree or in different ways. This is consistent with observations that low-level visual features are represented in high-level ventral visual cortex alongside features of higher complexity ^50–52^, and that categories are systematically related to category through differences in spatial frequency content, thus support classification ^53^.

We thus investigated the role of low-level features at different spatial frequencies in visual category representations in infants and adults. We filtered the stimulus material in spatial frequency in 100 bins spaced logarithmically between 0.1 and 30 cycle per degree (cpd) visual angle (for an example see **Figure 3E**). As expected, visual object categories were associated with different spatial frequency content in the images (**Figure S3E**) that allows category to be determined directly from the images (**Figure S3F**). Using RSA, we assessed the similarity between category representations and the spatially filtered images. We observed a significant relationship across all spatial frequencies (except at 0.18 cpd) in both infants and adults (**Figure 3G**), with stronger relationships for adults than infants above 1 cpd. This shows that category representations in infants and adults are differentiated by features across the spatial frequency spectrum, with a stronger role of higher spatial frequencies in adults.

Based on this result, we refined the deep neural network model based analysis with respect to spatial frequency. We compared the VGG model’s representation of the filtered images with infant and adult category representations (**Figure 3H**). For adults, the result revealed similar representations across all spatial frequencies as expected, with a peak at 3.06–5.42 cpd. In contrast, for infants, the similarity was restricted to spatial frequencies from 0.31 to 1.73 cpd, with a peak at 0.55–0.98 cpd. This is consistent with the shift in peaks in spatial sensitivity from low spatial frequency up to 1 cpd ^54,55^ to higher spatial frequency at 2–6 cpd ^56^. As expected from the previous analysis, we find significantly stronger correlations for higher spatial frequencies in adults than in infants.

Taken together, this reveals that in infants, category representations are in the format of low to intermediate complexity features at low spatial frequency and form a subset of the representations seen in adults. In contrast, in adults, category is discriminated by features at all levels of complexity and all spatial frequencies, with a higher reliance on high spatial frequencies.

Which processes may contribute to the emergence of high-complexity features in the developmental trajectory from infancy to adulthood? At this moment, we can only speculate. The results of the time-generalization analysis in conjunction suggest that local and far-reaching feedback processes from the frontal cortex might be involved ^57–59^. In more cognitive terms, linguistic and semantic processing is known to modulate visual processing in adults and might modulate visual representations ^60^.

Previous research investigating the format of infant visual category representations tested hypotheses one by one through experimental manipulation, for example, determining whether infants are sensitive to stimulus inversion ^27^ or tolerant to changes in viewing conditions ^61^. Instead, our approach allows the comparison of any number of hypotheses as captured in explicit, image-computable computational models to capture infant visual category representations in the future increasingly accurately. To speed up this process, we make the data publicly available.

Our findings further suggest constraints for artificial intelligence research. The biological brain inspired the engineering of deep learning models, but the models’ learning has remained biologically unrealistic ^12,49,62^ and is perceived as a major impediment to building better models. We suggest that models of human visual categorization striving for increased biological realism should follow a similar developmental trajectory of representations as described here.

### Spectral properties of visual category representations

Neural oscillations underlie the formation and communication of visual representations ^14,63–65^. Here we determined the spectral signature of visual category representations in infants as a first step toward describing their relationship to neural oscillations. For this, we resolved EEG data in distinct frequency bins from 2 to 30Hz and performed time-resolved visual category classification on each bin separately (**Figure 4A**). In infants (**Figure 4B**), we observed significant category classification accuracy in a specific cluster in the theta band with a peak at 4.63Hz (2.91–6.73Hz) and 400ms (160–580ms). This result reveals activity in the theta band as the spectral signature of visual category representations in infants. In contrast, in adults (**Figure 4C**), the cluster extended across the whole frequency range and time course investigated. It shows that the spectral signature of visual category representations in adults is broadband. The patterns of results did not depend on any particular category except on faces in infants (**Figure S4E,F**). Note that the observed differences in infants and adults are not a trivial consequence of differences in EEG power spectra pattern or category, as those were similarly broadband in both infants and adults and for all categories (**Figure S4A,B**). Further, the classification peaks do not map onto the power spectra peaks in terms of frequency and latency (**Figure S4C,D**). They are thus not a function of signal-to-noise ratio in the power spectrum. Finally, the difference between infants and adults is not due to higher inter-subject variability in spectral power patterns in adults, as different measures of variability were lower in adults than in infants (**Table S4D**).

**Figure 4.**
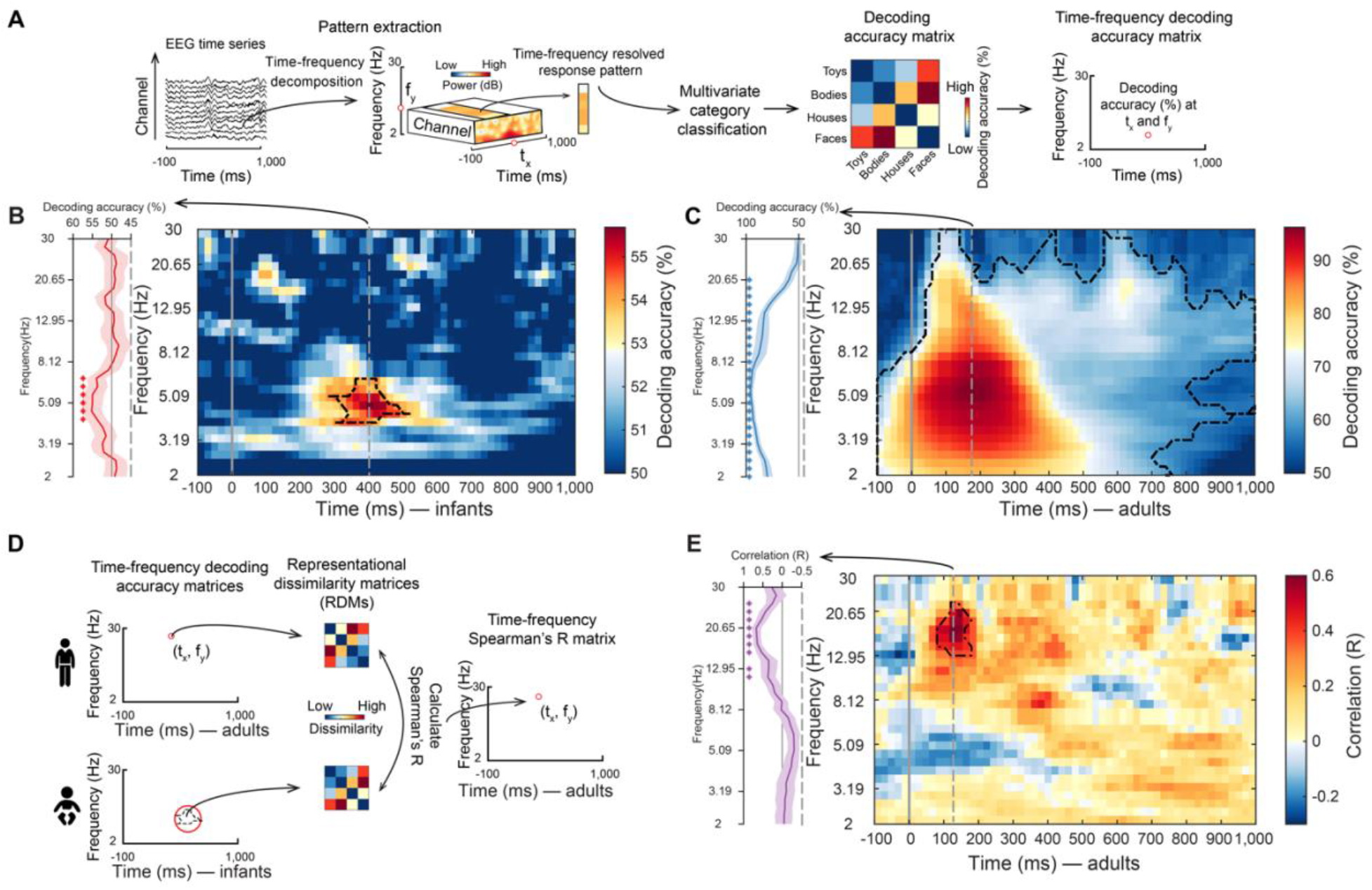
Spectral characterization of infant and adult category representations. **(A)** Category classification based on frequency-resolved EEG data. We first decomposed EEG data in time and frequency using *Morlet* wavelets for each trial and each channel, yielding a trial-wise representation of induced oscillatory power. We then conducted the time-resolved multivariate classification of category separately on each frequency bin. This yielded a 4 × 4 matrix of decoding accuracies at each time point and frequency bin, which we either averaged to obtain grand average category classification **(B,C)** or used as an RDM in RSA **(D,E)**. **(B,C)** Results of time- and frequency-resolved MVPA for infants **(B)** and adults **(C)**. Detailed statistical information is listed in **Table S4A**. **(D)** RSA procedure linking oscillation-based visual category representations in infants and adults. We first created a single aggregate infant oscillatory RDM by averaging decoding accuracy matrices based on the extent of the cluster in the infant data. We compared (Spearman’s *R*) this aggregate infant RDM to time- and frequency-resolved RDMs for each participant in the adult sample. This yielded a two-dimensional matrix indicating in which frequency range and when category representations are similar between infants and adults. **(E)** Similarity between infant theta-based category representations and adult category representations resolved in time and frequency. Detailed statistical information is listed in **Table S4B**. For **(B,C,E)**, for the single participant data see **GitHub Repository: Visualization**; the gray vertical lines indicate the onset of image presentation; line profiles of classification accuracy or correlation at the peak latency (gray vertical dashed lines) were shown on the left of the plots, respectively; black outlines indicate time point combinations with significant results (infants *n* = 40, adults *n* = 20, right-tailed permutation test, cluster-defining threshold *P* < .005, corrected significance level *P* < .05). See also **Figure S4** and **Table S4**.

The observed pattern of results is consistent with two alternative hypotheses about the relationship between the oscillatory basis of visual category representations in infants and adults. One hypothesis is that there is a direct match in frequency, suggesting that peak classification in infants and adults is at similar frequencies (i.e., at 4.63Hz and 5.59Hz, respectively). Another hypothesis is an upward shift across age, made plausible by the observations that brain rhythms increase in frequency during infant development ^66^.

To arbitrate between those hypotheses, we determined which frequency band and time points category representations in adults were similar to infant category representations identified in the theta band (**Figure 4D**). We extracted RDMs from the infant data at the cluster in the theta range. We used their average as a search template, comparing it to RMDs from the adult data for all time-point and frequency combinations. We found a cluster of significant correlations with a peak at 17.13Hz (9.78–20.65 Hz) at 120ms (120–360ms) (**Figure 4E**). This pattern of results was partly independent of category (except faces, **Figure S4G**), held across different data aggregation schemes and ways to assess the theta cluster (**Figure S4H**). Further, there was no relationship between signals at the (non-significant) peak in infant alpha/beta at 100ms and adult signals at any time-frequency combination (**Figure S4I,J**). It directly and specifically quantitatively demonstrates an upward shift in the spectral signature of neural activity supporting visual category representations from the theta range in infants to the alpha/beta range in adults. As expected from the investigation of the representational format above, the shared representations which shifted relied on low spatial frequency features (**Figure S3G,H**).

The shift observed is not a trivial consequence of differences in the peak latency and frequency of the power spectrum, which are similar in infants and adults (**Figure S4A–D**). Closer inspection of the classification results (**Figure 4B,C**) at peak latency reveals a similar peak around 4.63Hz– 5.59Hz (**Figure 4B,C**, line profiles), leaving open the possibility that to some degree the profile observed in infants might be a down-scaled and noisier version of the situation in adults, and predicting shared representations across age group in the theta band. Instead, the shared representations are present only in the alpha/beta band (**Figure 4E**, line profile), demonstrating a clear dissociation from overall signal strength.

One interpretation of these findings is that in infants, neural networks for learning and memory associated with the theta rhythm contribute to the formation of category representations ^67,68^, whereas in adults, equivalent category representations are processed quickly in fully developed semantic networks associated with the alpha/beta rhythms ^69,70^. Alternatively, the frequency shift might be due to more efficient axonal transmission as a result of improved myelination ^40^ that may enable higher neural oscillations to emerge in the same neural circuits, consistent with increases in the prevalent frequency in the EEG across development ^66^. On this account, our finding suggests a novel general developmental trajectory for neural communication channels in the human brain: specific information-processing mechanisms working at low frequency in infants are shifted up in frequency gradually across development, and the size of this shift depends on the differences in neural circuit myelination. However, we note that power in a frequency does by itself indicate oscillations in that frequency, and further research is needed to establish this firmly, e.g., by distinguishing aperiodic from periodic components ^71,72^. Similarly, we do not observe a one-to-one mapping between shared representations revealed by time-resolved analysis (**Figure 2B**) and time-frequency resolved analysis (**Figure 4E**). Instead, we expect the relationship to be akin to the complex relationship between neural oscillations and evoked responses ^73,74^. Further research is needed to resolve to which degree they depend on distinct or shared neural phenomena.

### The nature and developmental trajectory of infant to adult visual category representations

In sum, our results reveal the nature and developmental trajectory of the infant to adult visual category representations, from infancy to adulthood. Temporal dynamics change from slowly to quickly emerging in time, the format from visual features up to intermediate complexity to features of high complexity, and the oscillatory signature from the theta to the alpha/beta frequency. These results provide insight into visual category representations that underlie the development of fast and efficient visual categorizations skills in humans. They also further reveal how cortical information transmission channels change in human development and demonstrate the power of advanced multivariate analysis techniques in infant EEG research for developmental cognitive science.

## ACKNOWLEDGMENTS

R.M.C is supported by Deutsche Forschungsgemeinschaft (DFG) grants (CI241/1-1, CI241/3-1, CI241/7-1) and by a European Research Council Starting Grant (ERC-2018-StG). S.X. is supported by a scholarship by the Chinese Scholarship Council. M.K. and S.H. were supported by the DFG and FWF jointly (grant numbers: KO 6028/1-1; I 4332-B), and S.H. also by the Max Planck Society. Computing resources were provided by the high-performance computing facilities at ZEDAT, Freie Universitaet Berlin.

## AUTHOR CONTRIBUTIONS

Conceptualization: RC, SX, MK, SH, BT. Methodology: SX, RC, MK. Software: SX, MK. Formal analysis: SX, MM. Investigation: EK, CK, Resources: SX, RC, MK. Data curation: SX. Writing - original draft preparation: SX, RC. Writing - review and editing: RC, SX, MK, SH, BT. Visualization: SX. Supervision: RC, MK. Project administrations: RC, SH, MK. Funding acquisition: RC, SH and MK.

## STAR METHODS TEXT

### KEY RESOURCES TABLE

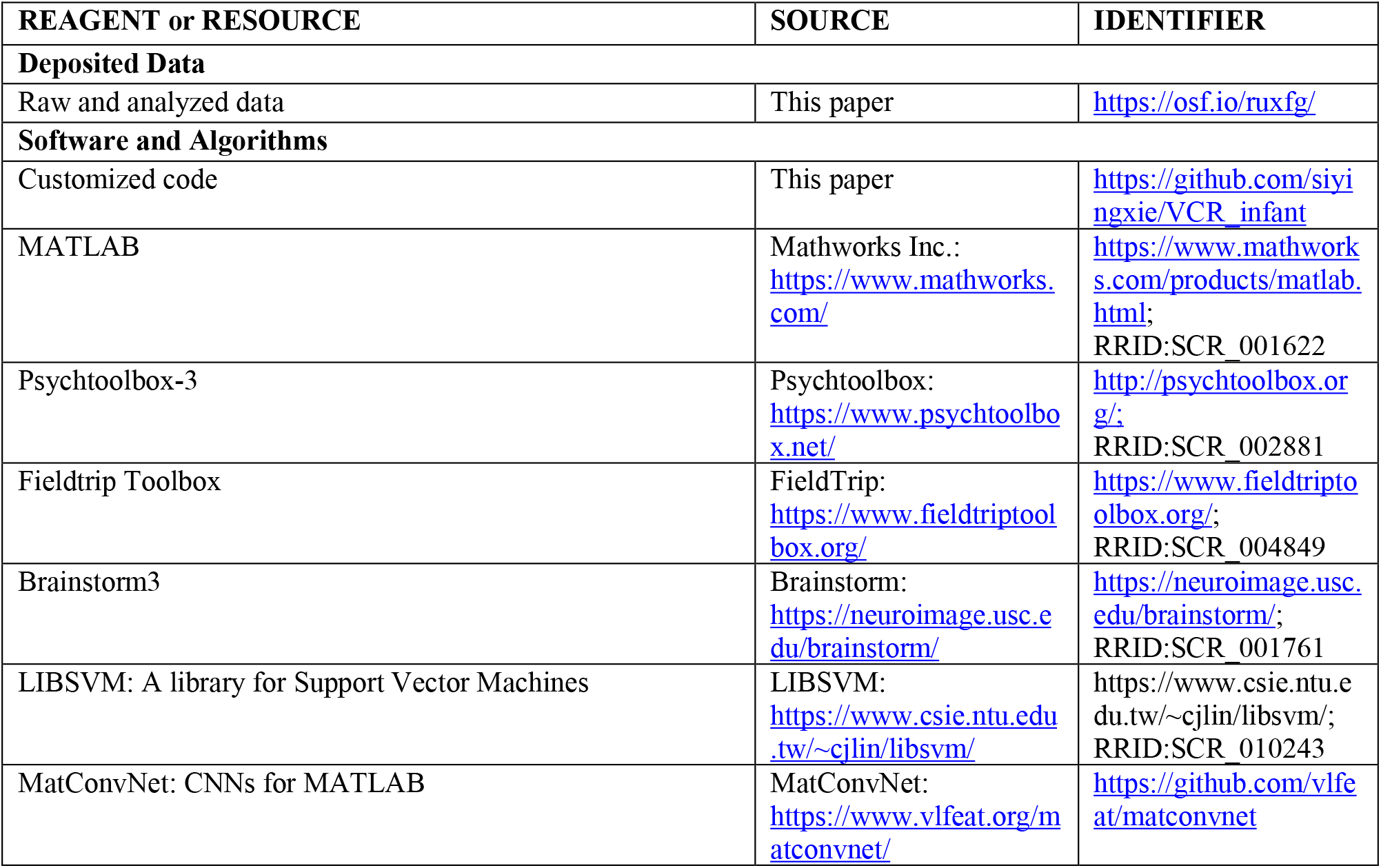

### RESOURCE AVAILABILITY

#### Lead contact

Further information and requests for the resources should be directed to and will be fulfilled by the lead contact, Radoslaw M. Cichy.

#### Materials availability

This study did not generate new unique reagents.

#### Data and code availability

Raw and processed data have been deposited at *OSF* and are publicly available as of the date of publication. DOI is listed in the key resources table.

All customized codes have been deposited at *GitHub* and are publicly available as of the date of publication. DOI is listed in the key resources table.

### EXPERIMENTAL MODEL AND SUBJECT DETAILS

Two independent pools of participants took part in this study: 6–8 months old infants and young adults. We chose this infant age group based on extensive evidence from behavioral and electrophysiological work showing that infants discriminate between various basic level visual categories by this age, whereas in younger infants, category discrimination is less stable and relies more on the chosen paradigm and the specific categories assessed ^26,29–31^. The infant sample was assessed at the Max Planck Institute for Human Cognitive and Brain Sciences in Leipzig, Germany. It comprised 48 participants, of which 8 were excluded due to insufficient data, yielding a final sample analyzed of 40 infant participants (gender: 19 female, age: mean ± SD: 214.9 ± 14.76 days). The adult sample was assessed at the Freie Universität Berlin, Germany. It comprised 20 participants of which none was excluded (gender: 11 female, age: mean ± SD: 26.1 ± 3.81 years). Caregivers of all infants and all adult participants gave written informed consent. The study was conducted according to the Declaration of Helsinki and the infant and adult protocols were approved by the respective local ethic committees.

### METHOD DETAILS

#### Stimuli

The stimulus set consisted of 32 object images in each of the four categories included: houses, toys, faces, and bodies. We chose those categories for four reasons: (1) they are highly familiar to infants, and infants encounter them in everyday life; (2) faces ^75–78^, toys ^3^, bodies ^79,80^, and houses ^81,82^ (as large objects that define scenes) have been used in previous infant research making our results in principle comparable; (3) they have well-described and distinct neural signatures in adults ^83^; and (4) a recent fMRI study showed distinct neural signatures for faces, objects, and scenes in infants, too ^20^. This yielded a total set of 4 × 32 = 128 object images. All object images were cut-out from color photographs. All images are available in the **OSF repository**. We analyzed the data at the level of category.

#### Experimental procedure in the infant sample

In the infant experiment, participants were presented with 272 trials divided into four blocks. Each block had the same basic structure. At the beginning of each block four stimuli (one stimulus per object category) were separately presented three times in randomized order. Thereafter participants were presented with a random sequence of images comprising the same four stimuli seven more times, intermixed with 28 other images (seven images per category) presented only once. This experimental design was chosen because it allows assessing the effect of object image repetition of infant brain responses, but this question is orthogonal to the ones pursued here and will be reported separately.

Each trial consisted of a fixation dot presented for a variable duration of 700–900ms, followed by a stimulus presented for 2,000ms at the center of the screen (**Figure S1A**). To capture the attention of the infants and direct their gaze to the screen we implemented two measures. First, we presented a yellow duck image and duck sound for 1,000ms at the beginning of each block and thereafter every 10 trials. Second, each stimulus was presented together with one of ten arbitrary sounds that were assigned randomly at each trial.

During the assessment, infants sat on their care giver’s lap at a viewing distance of about 80 cm from a 17-in. CRT screen. The object images were presented at the center of the screen, subtending a visual angle of approximately 5.0°. To monitor infants’ gaze, we recorded videos of infants’ faces throughout the experiment.

#### Experimental procedure in the adult sample

We adapted the experimental design for the adult sample. In short, all images were shown equally often, with higher number of repetitions, at shorter presentation times and higher presentation rates that in the infant study (**Figure S1B**).

The first 3 participants were presented with 1,280 trials divided into 5 runs. In each run each object image was presented twice. The other 17 participants were presented with 3,840 trials divided into 10 runs. In each run each object image was presented three times. In each run, images were presented in random order, and runs were separated by breaks that were self-paced by the participants.

Each trial consisted of the presentation of a fixation cross with a variable duration of 600–800ms, followed by a stimulus presentation for 500ms. Stimuli were presented at the center of the screen at a visual angle of approximately 7.0°.

Participants were instructed to keep fixation on the center of the screen throughout the experiment. To ensure that participants attended to the stimuli and to avoid contamination of the relevant recording times with blink artefacts, participants were instructed to press a button and blink their eyes in response to a paper clip image that was shown randomly every 4 to 6 trials (average 5 trials). Paper clip trials were excluded from all further analysis.

#### EEG acquisition and preprocessing

##### Infant sample

EEG data for the infant sample were recorded in a shielded room using 30 Ag/AgCl ring electrodes and a TMSi 32-channel REFA amplifier at a sampling rate of 500 Hz. Electrodes were placed according to the standard 10-20 system. Electrodes V+Fp2 and V− recorded the vertical electrooculogram (VEOG), and electrodes H-F9 and HF+10 recorded the horizontal electrooculogram (HEOG), Cz served as the online reference. We conducted preprocessing using the Fieldtrip toolbox ^84^. The continuous EEG data was segmented for each trial into epochs. For subsequent time-resolved multivariate analysis we extracted the epoch from −200ms to +1,000ms with respect to image onset. For analysis that was additionally resolved in frequency we used longer epochs to allow better estimation at lower frequencies from −500ms to +1,000ms.

We removed all trials during which participants did not gaze at the screen for 1,000ms after stimulus onset as assessed by visual inspection of the video recordings. In this reduced trial set (mean ± SD: 139.6 ± 47.76 trials) we removed noisy channels (mean ± SD: 1.25 ± 1.32) and replaced them by interpolated data from adjacent electrodes. We further conducted independent component analysis (ICA) and removed components related to eye-movement and muscle artifacts as identified by visual inspection.

##### Adult sample

EEG data for the adult sample were recorded using an EASYCAP 64-channel system and a Brainvision actiCHamp amplifier at a sampling rate of 1,000Hz. Data were filtered online between 0.3 and 100 Hz. Electrodes were placed according to the standard 10-10 system. Electrode Fz served as the online reference. We conducted preprocessing using the Brainstorm 3 toolbox ^85^. Up to two noisy channels were removed for each participant as identified by visual inspection. We conducted ICA to identify and remove eye-movement and muscle artifact components by visual inspection of independent components. The continuous EEG data were then segmented for each trial into epochs from −200ms to +1,000ms (for time-resolved analysis) and from −500ms to +1,000ms (for time- and frequency resolved analysis).

#### EEG time-frequency decomposition

We decomposed the EEG time series into frequency-specific components by convolving the data with complex *Morlet* wavelets separately for each trial and sensor. We performed decomposition based on single trials so that the decomposed activity reflects stimulus-locked evoked responses and induced responses ^73^. The wavelets had a constant length of 2,600ms and were logarithmically spaced in 30 frequency bins between 2Hz and 30Hz. We obtained the absolute power values for each time point and frequency bin by taking the square root of the resulting time-frequency coefficients. We normalized these power values to reflect relative changes (expressed in dB) with respect to the pre-stimulus baseline (−300ms to −100ms with respect to stimulus onset). We downsampled the time-frequency representations to a temporal resolution of 50 Hz (by averaging data in 20ms-bins) to increase the signal-to-noise ratio of subsequent analyses. This yielded for each trial a power value for each time point and frequency bin.

#### Multivariate classification of visual category from EEG data

To characterize the temporal dynamics with which visual category representations emerge in infant and adult brains we conducted multivariate EEG classification using linear support vector machines (SVMs). We analyzed the infant and the adult data set separately and equivalently.

We conducted two common variants of multivariate EEG classification: time-resolved EEG analysis ^9,37^ and time-generalization analysis ^39^. We conducted the analysis separately on the adult and infant sample, and separately for each participant. All analyses employed binary c-support vector classification (C-SVC) with a linear kernel as implemented in the LIBSVM toolbox ^86^. The details of the time-resolved and the time-generalization analysis are as follows.

##### Time-resolved classification

We used time-resolved multivariate pattern analysis on EEG data (**Figure 1B**) to determine the time course with which visual category representations emerge in infant and adult brains. For each time point of the EEG epoch (from −200ms to +1,000ms), we extracted trial-specific EEG channel activations (i.e., 25 in infants and 63 in adults) and arranged them into pattern vectors for each of the four category conditions (i.e., face, house, body, and toy) of the stimulus set. To increase the signal-to-noise ratio (SNR), we randomly assigned raw trials into four bins of approximately equal size each and averaged them into four pseudo-trials. We used a leave-one-pseudo-trial-out cross validated classification approach. We trained the SVM classifier to pairwise decode any two conditions using three of the four pseudo-trials for training. We used the fourth left-out pseudo-trial for testing, yielding classification accuracy (chance level 50%) as a result. The procedure was repeated 100 times, each time with a new random assignment of trials to pseudo-trials. The resulting decoding accuracy was averaged across repetitions and assigned to a decoding accuracy matrix of size 4 × 4, with rows and columns indexed by the conditions classified. The matrix is symmetric across the diagonal, with the diagonal undefined. This procedure yielded one decoding matrix for every time point.

##### Time-frequency resolved classification

In addition to classifying visual category from broadband responses (i.e., single trial raw unfiltered waveforms), we classified object categories from oscillatory responses. This analysis followed the same rationale as the classification analysis described above, with the only difference that classification was conducted on power value patterns instead of raw activation value patterns. The analysis was conducted separately for each frequency bin separately. This resulted in a decoding accuracy matrix of size 4 × 4 as defined above for every time point and every frequency bin.

##### Time generalization analysis

We used time-generalization classification analysis ^39^ to determine how visual representations emerging at different time points during the dynamics of visual perception relate to each other. For time and memory efficiency, we down-sampled the EEG data to a sampling rate of 50 Hz by averaging the raw EEG data in 20ms bins. The procedure was equivalent to the time-resolved classification analysis with the only difference that classifiers trained on data from a particular time point were not only tested on left out data from the same time point, but iteratively on data from the same and all other time points. The idea is that successful classifier generalization across time points indicates similarity of visual representations over time. This analysis yielded thus a size 4 × 4 decoding accuracy matrix indexed in rows and columns by the conditions compared for all time point combinations from –200 to +1,000ms. We averaged the entries of the decoding accuracy matrix at each time point, yielding a temporal generalization matrix indexed in rows and columns by training and testing time.

##### Sensor-space searchlight analysis

We performed a sensor-space searchlight analysis ^87,88^ to localize in EEG channel space which channels contributed to the classification of category. For each EEG channel we defined a neighborhood as a sphere of the 10 (for adults) or 5 (for infants) closest EEG channels. For each EEG channel we then performed time-resolved category classification analysis, limiting data entering the analysis to its neighboring channels. Averaging across all pairwise category classifications yielded one decoding accuracy for each time point and for each EEG channel. We further averaged the results in 200ms bins, yielding a single EEG channel searchlight map of grand average decoding accuracy for each time bin.

#### Computing spatial-frequency specific versions of the stimulus set

To assess the role of spatial frequency on visual object categorizations, we decomposed the stimulus set in terms of spatial frequency. For this, we first used the Fourier transform to transform each image into the frequency domain. We then defined a set of 100 Butterworth band-pass filters (complex higher order filters with a roll-off response rate of 5) logarithmically spaced between 0.1 and 30 cycle per degree (cpd) visual angle. We applied each band-pass filter to the frequency representation of each image, yielding 100 band-pass filtered versions of each image in the frequency domain. We combined the resulting power values of each image in the frequency domain together with the image’s original phase information to compute the corresponding frequency-filtered images using the inverse Fourier transform. This procedure resulted in 100 sets of the stimulus set, band-pass filtered between 0.1 and 30 cpd.

#### Comparing visual representations in infants and adults

We determine whether infants and adults have similar visual category representations using representational similarity analysis (RSA) ^89,90^. The idea is that infants and adults share representations of category if they treat the same categories as similar or dissimilar. We determined this in a two-step process. In a first step, for each age group independently condition-specific multivariate activity patterns (adults: 63 electrodes; infants: 25 electrodes) were compared for dissimilarity. Dissimilarity was determined for all pairwise combinations of conditions, and dissimilarity values were aggregated in so-called representational dissimilarity matrices (RDMs) indexed in rows and columns by the conditions compared (here: 4 × 4 RDMs indexed by the 4 object categories). RDMs thus provide a statistical summary of the similarity and thus representational relations between visual category representations. The RDMs gained from the infant and adult sensor space separately have the same definition and dimensionality and are thus directly comparable. Thus, in a second step, the infant RDM and the adult RDMs are related to each other by determining their similarity.

We applied RSA to two different types of data: evoked responses (i.e., recorded voltage signals) and oscillatory responses (i.e., spectral power). In both cases we re-used the results of the classification analysis described above for the definition of RDMs. Classification accuracy can be interpreted as a dissimilarity measure on the assumption that the more dissimilar activation patterns are for two conditions, the easier they are to classify ^9,91^. We detail the different RSA procedures below. To reduce visual complexity of the analysis we subsampled the results of the classification analysis by binning them in 10ms bins.

##### Relating visual category representations in infants and adults based on raw broadband time courses

We investigated whether infants and adults share common visual representations based on broadband responses. As visual representations in adults and infants likely emerge with different time courses, we related their visual representations in a representational similarity time-generalization analysis. As RDMs we used time-point specific decoding accuracy matrices (**Figure 2A**). We first averaged infant RDMs across all participants to increase SNR, resulting in one average infant RDM per time point. We then correlated (Spearman’s *R*) the average infant RDM to each adult (*n* = 20) RDM across all time point combinations. This yielded 20 correlation matrices, indexed in rows and columns by the time points compared (rows: infant time; columns: adult time), indicating when infants and adults share category representations.

##### Relating visual category representations in infants and adults based on frequency-specific power time courses

We investigated whether infants and adults share visual representations in particular frequency bands. As RDMs we used decoding accuracy matrices from the classification analysis based on time-frequency resolved power values. As in this analysis we could neither assume similar time courses, nor similar roles for particular frequencies across infants and adults, we related infant and adult representations in a time-and-frequency-generalization analysis. To do this, we first defined a single aggregate infant RDM by averaging decoding accuracy matrices based on the extent (time and frequency) of the significant cluster in the infant data (**Figure 4B**) alone. To increase signal-to-noise we only included RDMs whose average across entries in single participants was greater than or equal to 50% decoding accuracy. Note that this criterion is orthogonal to the hypotheses tested and thus does not bias the analysis. We compared (Spearman’s *R*) this single aggregate infant RDM to time- and frequency-resolved RDMs for each participant of the adult sample (*n* = 20), separately for each frequency and time point. This yielded 20 correlation matrices, with rows representing time points and columns representing frequency bins, indicating when and at which frequency infants and adults share category representations.

##### Relating visual representations in infants and adults to computational models

To characterize the format of visual category we related neural representations in infants and results to different computational models using RSA. We constructed model RDMs from computational models (**Figure 3A**) that represent visual information in different formats. We considered two types of visual computational models: a Gabor wavelet pyramid as a model of low-level feature representations ^47,92^, and the VGG-19 ^48^ deep convolutional neural network (DNN) trained to categorize object images. Deep neural networks process visual information along a hierarchy of increasing complexity from low to high ^93,94^ that has been shown to match the processing hierarchy of the human brain ^46,95,96^ and predict human and non-human primate brain activity better than other model class ^12,97–99^.

To construct model RDMs we first ran all visual stimuli in the study (i.e., 128 object images) through the models and extracted their activation values. More specifically, for the Gabor model we extracted a single set of model responses for Gabor wavelets differing in size, position, orientation, spatial frequency and phase. For the DNN we used the MatConvNet toolbox ^100^ to extract model neuron activation values from the rectified linear units (Relu) for each layer. We z-transformed activation values across stimuli for each stage/layer separately and averaged the transformed values across the 32 stimuli belonging to each of the four categories (i.e., face, body, house and toy), resulting in four category-specific activation values. We formed the patterns into vectors and computed the dissimilarity (1 – Pearson’s *R*) between all pairwise combinations of the four category activation vectors, resulting in a 4 × 4 RDM for each DNN layer of each DNN separately, and one model RDM for the Gabor feature model.

To construct spatial frequency-specific DNN model RDMs, we used an equivalent procedure with the difference that we ran band-pass filtered images through the DNN model separately for each band-pass defined. This resulted in 4 × 4 RDM for each spatial frequency band and DNN layer of the DNN separately.

To construct spatial-frequency specific image-based RDMs, we did not run the images through a model, but the procedure was otherwise equivalent. We directly averaged the pixel values of the filtered images across the 32 stimuli belonging to each of the four categories (i.e., face, body, house, and toy), resulting in four category-specific activation values. We formed the patterns into vectors and computed the dissimilarity (1 - Pearson’s *R*) between all four category activation vectors pairwise combinations, resulting in a 4 × 4 RDM for each frequency band.

To construct neural RDMs that capture category representations well we averaged decoding accuracy matrices from time-resolved category classification (**Figure 1C,D**) in the 95%confidence intervals around peak latency in time-resolved category classification. Our rationale was that peak latency is the time point when categories were linearly best separable and thus their representations most explicit ^101^. To increase signal-to-noise we only included RDMs whose average across entries in single participants was greater than or equal to 50% decoding accuracy. Note that this criterion is orthogonal to the hypotheses tested and thus unbiased. This yielded a single neural RDM for every infant and adult participant. We then related infant and adult neural RDMs to model RDMs using Spearman’s *R*, yielding a single correlation value for each model RDM and participant.

To allow assessing the models’ predictivity with respect to the noise in the data we calculated an upper and lower bound for the noise ceiling ^102^, that is the predictions a perfect model may reach given the noise in the data. This procedure was conducted separately for the infant and the adult sample. To estimate the upper bound we correlated (Spearman’s *R*) each participant’s neural RDM with the mean neural RDM across all participants. To estimate the lower bound we correlated (Spearman’s *R*) each participant’s neural RDM with the mean neural RDM excluding that participant iteratively for all participants. We averaged the results, yielding estimates of the lower and upper noise ceiling for infants and adults.

To reveal whether infants, adults, and the DNN share common representations, we applied variance partitioning using a general linear model (GLM). The procedure was as follows. We first computed two GLMs between the DNN model RDM (i.e., observation) and the infant and adult RDMs (i.e., main regressor), respectively. This revealed the total variance that the model and each age group shared. We then computed two additional GLMs, adding the other age group’s average RDM to the model (i.e., there are two main regressors). From the additional RDMs, we obtained the unique variance of infant and adult RDMs, which was the difference in explained variance after infant and adult RDMs were reduced from the models. From the results of those two types of GLMs, we computed the shared variance that resulted from subtracting the unique variance from the total variance. We applied this analysis for network layers to which infants and adults showed a significant relationship in the correlation analysis, which are layers 3 and 4.

### QUANTIFICATION AND STATISTICAL ANALYSIS

We used non-parametric statistical inference for random-effects inference to avoid assumptions about the distribution of the data ^103,104^. We used permutation tests for cluster-size inference, and bootstrap tests for confidence intervals on maxima, cluster onset/offset, and peak-to-peak latency differences. The sample size (*n*) for infants was 40 and for adults 20. Tests were either two- or right-tailed and are indicated for each result separately.

#### Permutation tests

We tested the statistic of interest (i.e., mean decoding accuracy or correlation coefficient in RSA across participants) using sign permutation tests. The null hypothesis was that the statistic of interest was equal to chance (i.e., 50% decoding accuracy, a Spearman’s *R* of 0). Under the null hypothesis, we could permute the category labels of the EEG data, which effectively corresponds to a sign permutation test that randomly multiplies participant-specific data with +1 or –1. For each permutation sample, we recomputed the statistic of interest. Repeating this permutation procedure 10,000 times, we obtained an empirical distribution of the data. We converted the original statistic (i.e., correlation coefficient, the decoding time courses, time-time matrices of correlation coefficients or decoding accuracies, and time-frequency decoding matrices) into *P*-values (correlation coefficients), 1-dimensional (time courses), or 2-dimensional (time-generalization or time-frequency) *P*-value matrices.

We controlled the family-wise error rate using cluster-size inference. We first thresholded *P*-value time courses or maps at *P* < .005 (cluster-definition threshold) to define supra-threshold clusters by contiguity. These supra-threshold clusters were reported significant only if the size exceeded a threshold, estimated as follows: the previously computed permutation samples were also converted to *P*-value time courses/matrices and also thresholded to define resampled versions of supra-threshold clusters. These clusters were used to construct an empirical distribution of maximum cluster size and estimate a threshold of 5% of the right tail of this distribution (i.e., the corrected *P*-values is *P* < .05).

#### Bootstrap tests

We calculated 95% confidence intervals for the onsets of the first significant cluster and the peak latency of the observed effects. To achieve this, we created 1,000 bootstrapped samples by sampling the participants with replacement. For each bootstrap sample, we determined the peak latency as well as onsets of the first significant cluster and the offset of the last significant cluster. This resulted in empirical distributions of peak, onset and offset latencies on which we determined 95% confidence intervals.

To calculate confidence intervals on mean peak-to-peak latency differences, we created 1,000 bootstrapped samples by sampling the participant-specific latencies with replacement. This yielded an empirical distribution of mean peak-to-peak latencies. If the 95% confidence interval did not include 0, we rejected the null hypothesis of no peak-to-peak latency differences. The threshold *P* < .05 was corrected for multiple comparisons whenever appropriate using FDR correction.

---

**Figure S1.**
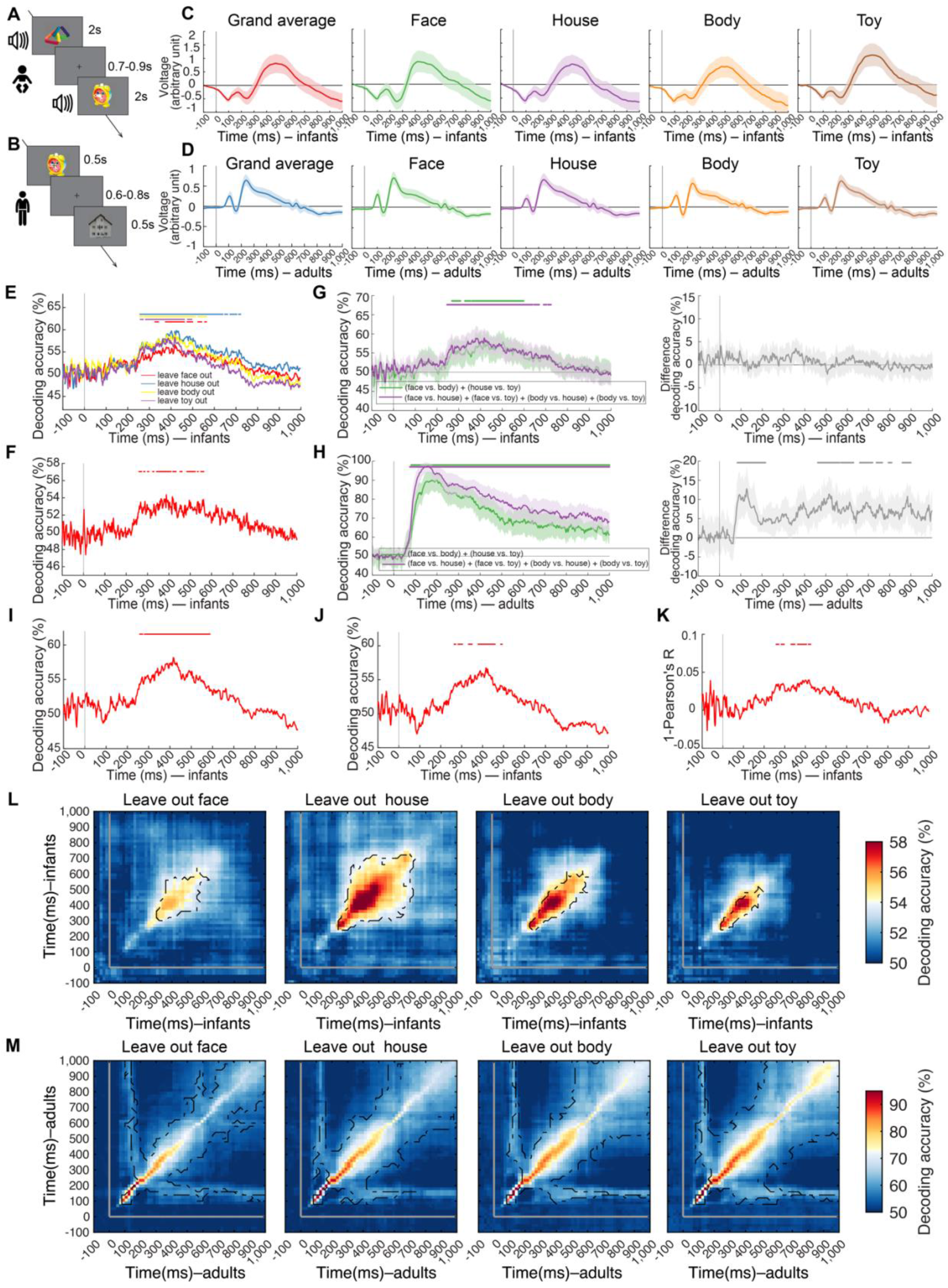
Analysis of Category Representations in Time for Infants and Adults, related to Figure 1 and STAR Methods. **(A,B)** Experimental paradigms. For the infant sample **(A)**, on each trial an object image was presented for 2s together with a randomly assigned 10 arbitrary sounds, followed by a variable inter-stimulus-interval (ISI) of 0.7s to 0.9s; for adult sample **(B)**, presentation parameters were adapted to increase stimulus repetitions and thus the signal-to-noise ratio. On each trial an object image was presented for 0.5s followed by a variable ISI of 0.6s to 0.8s. **(C,D)** Visualization of event related potentials (ERPs). We averaged baseline-normalized (mean removal and division by standard deviation of the baseline activity) trials (electrodes: O1, O2, P7, P8) across category (i.e., grand average) or by category (i.e., face, house, body and toy). The ERPs were plotted in **(C)** for infants and in **(D)** for adults. In each case, the reported ERPs were either the grand average across all categories or averaged for each category separately. P100 peak latencies were slightly but significantly delayed in infants (i.e., 22–68ms difference). Overall peak latency differences were moderate (i.e., 98–242ms difference). For detailed peak latencies and statistical information, see **Table S1F**. For visualization, results were smoothed within a 20-ms sliding window. Shaded areas above and below curves indicate 95% confidence intervals determined by bootstrapping the participant sample (1,000 iterations). The gray vertical line indicates the onset of image presentation. **(E,F)** Time-resolved multivariate analysis in infants do not depend on any particular object or category. To determine whether the pattern of results depended strongly on any one category, we averaged category classification results leaving out one of the four categories in turn. The results in **(E)** are equivalent to **Figure 1C**. To determine whether the time-resolved results depended strongly on any one object in the stimulus set, we performed category decoding using a leave-one-object-out cross-validation scheme. In detail, we trained the classifier using data from 31 of the 32 images of a category (e.g., 31 faces vs. 31 toys), and tested it on the left-out images (e.g., the 32nd face and toy). The results in **(F)** are equivalent to **Figure 1C**. Detailed statistical information for **(E,F)** is listed in **Table S1B**. **(G,H)** Results of time-resolved multivariate analysis held equally within and across the animacy distinction in both infants and adults. We grouped pairwise category classification results by the animacy of the categories classified. In detail, we averaged the results from 1) classifying within the animate (face vs. body) and within the inanimate (house vs. toy) division together (green curves); 2) classifying across the animate and the inanimate division (face vs. house, face vs. toy, body vs. house and body vs. toy) together (magenta curves). In infants **(G)**, we observed statistically significant results for classification within and across the animacy division separately (left panel) but the difference was not significant (right panel). In contrast and as expected (Carlson et al., 2013; Cichy et al., 2014); in adults **(H)**, we observed statistically significant results for classification both within and across the animacy division (left panel) and significantly higher results across the animacy division (right panel). It shows that infants represent finer distinctions than the basic animate vs. inanimate division and suggests that visual category representations do partly reflect the animacy division in adults, but not yet in our infant sample. The shaded margin on curves indicates 95% confidence intervals of decoding accuracy. **(I–K)** Results of time-resolved multivariate analysis in infants emerged equivalently for alternative analysis schemes. Category classification results did not strongly depend on particular preprocessing steps: results without **(I)** or with **(J)** univariate noise normalization (i.e. each trial and channel separately was z-transformed based on baseline activity from −200 to 0ms). Category differentiation showed an equivalent result pattern **(K)** when comparing category-specific EEG response patterns using 1-Pearson’s R rather than decoding accuracy as a measure. Detailed statistical information for **(I–K)** is listed in **Table S1C,D**. **(L,M)** Results of time generalization analyses showed robustness in both infants and adults when leaving out each one of the four categories To determine whether the pattern of results depended strongly on any one category, we averaged category classification results leaving out one of the four categories in turn. The results in **(L,M)** are equivalent to **Figure 1G,H**, respectively. We observe a similar result pattern as in the main analysis in each case. Notes: The gray lines on plots indicate image onset. Rows of asterisks in **(E–K)** indicate significant time points. Black outlines in **(L,M)** indicate time point combinations with significant results (infant *n* = 40, adult *n* = 20, right-tailed sign permutation tests, cluster-defining threshold *P* < .005, corrected significance level *P* < .05).

**Figure S2.**
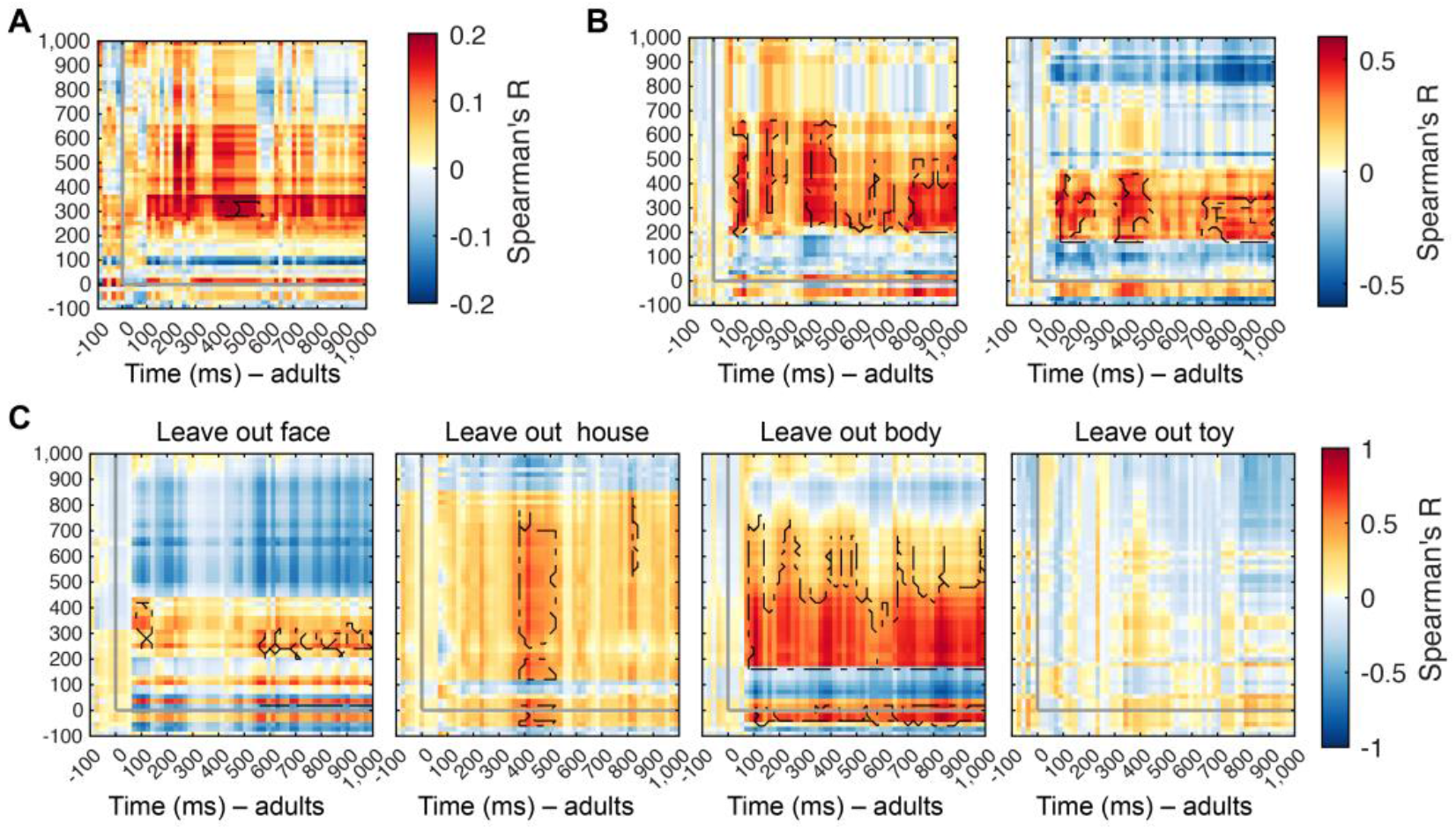
Analysis of Shared Category Representations between Infants and Adults, related to Figure 2. **(A,B)** Results of time-generalization RSA was similarly achieved for alternative processing and data aggregation choices. We compared RDMs (Spearman’s *R*) in infants and adults averaging over adult participants, rather than infant participants and the result was shown in **(A)**. Detailed statistical information is listed in **Table S2B**. We used 1 - Pearson’s *R* in (**B**, left) and Euclidean distance in (**B**, right) as a dissimilarity measure between brain patterns rather than decoding accuracy. Detailed statistical information is listed in **Table S2C,D**. **(C)** Results of time-generalization RSA did not depend on any particular category except on toys. To determine whether the pattern of results depended strongly on any one category, we left out one of the four categories in turn for RSA. We observe evidence for shared visual category representations in all analyses except the leave-toy-out analysis. Notes: The gray lines on plots indicate image onset. Black outlines indicate time point combinations with significant results (right-tailed sign permutation tests, cluster-defining threshold *P* < .005, corrected significance level *P* < .05).

**Figure S3.**
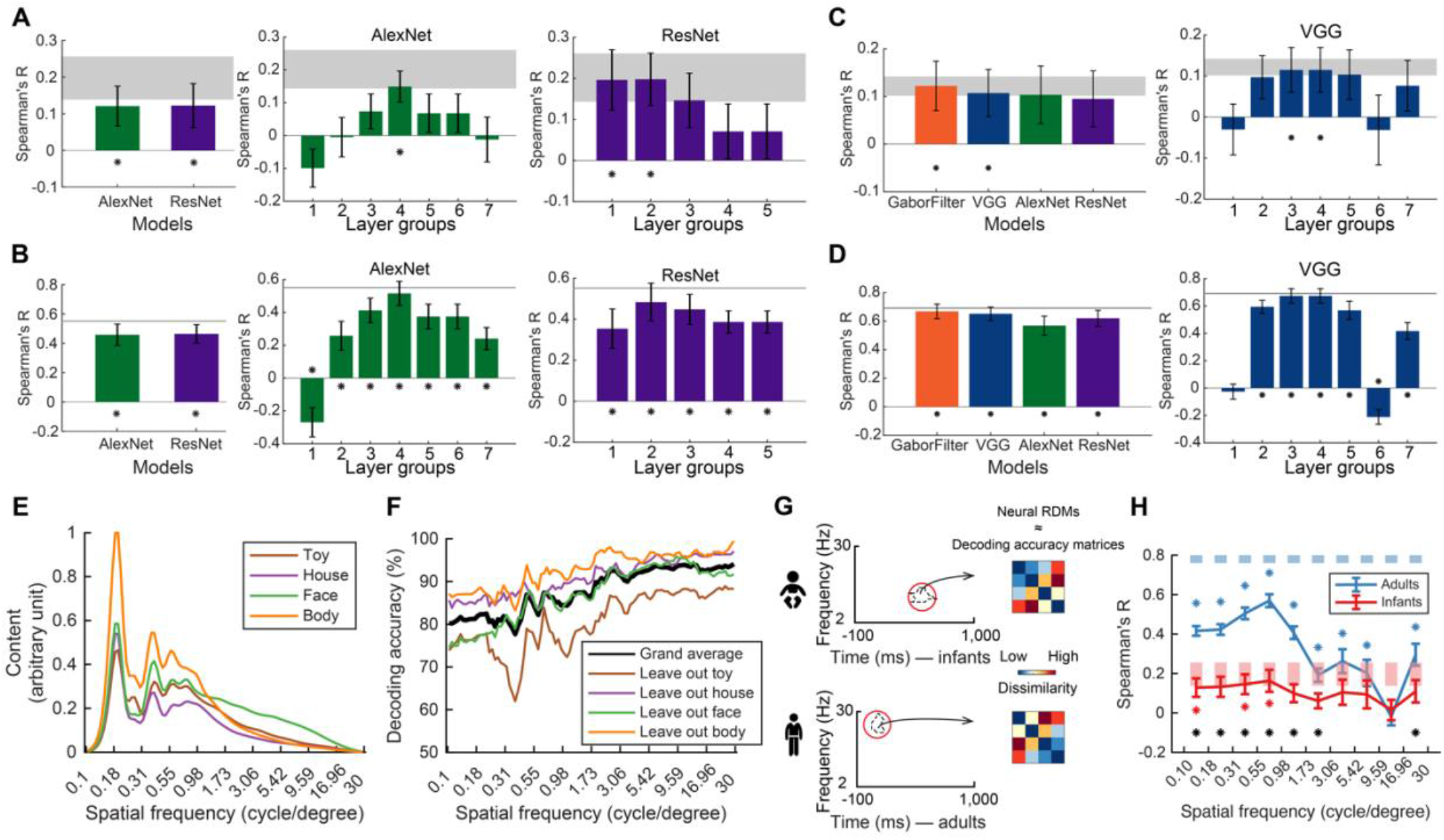
Analysis of the Format of Category Representations in Infants and Adults, related to Figure 3. **(A,B)** We assessed the format of category representations in infants and adults with relation to various DNNs (i.e., AlexNet and ResNet-50). Results for infants **(A)** and adults **(B)** at the whole-model level and DNN layer level. Statistical details (i.e., correlations and *P*-values) are in **Table S3A**. **(C,D)** We assessed the format of category representations using the significant shared representations, as shown in Figure 2B. For this, we defined neural RDMs as the average of decoding accuracy RDMs from time-resolved category classification in the temporal extent of the cluster indicating shared representations. Results for infants **(C)** and adults **(D)** at the whole-model level and at the DNN layer level for VGG models. Statistical details (i.e., correlations and P-values) are in **Table S3B**. **(E)** Spatial frequency spectrum in image categories. The panel showed the mean spatial frequency power in images for objects, ordered by category. Values are normalized such that the maximum value across categories is set to 1. The procedure was as follows. All images were filtered into one hundred bins logarithmically spaced between 0.1 and 30 cycles per degree visual angle using Butterworth band-pass filters. To compute image-specific spatial frequency power, each original image was first Fourier transformed into the frequency domain, multiplied by a 2-D frequency band-pass filter, and the power value was extracted as the spatial frequency content at that frequency bin. **(F)** Results of category classification from the spatial-frequency-specific power values of the images, averaged across all comparisons (grand average) and for each analysis leaving one category out. **(G)** Spatial-frequency-specific features in representations shared between adults and infants. For this, we extracted RDMs from the time-frequency classification analysis in the theta cluster in infants, and RDMs from the time-frequency classification analysis in adults at the location of the cluster indicating shared representations between infants and adults. We compared each in turn to spatial-frequency specific RDMs of the DNN. **(H)** Results indicate that adults represent features at all spatial frequencies, whereas infants do so only in low spatial frequencies. The conjunction of those results reveals that the features shared between infants and adults are in low spatial frequencies. Notes: For **(A–D,H)**, error bars represent standard errors of the mean. For **(A–D)**, asterisks indicate statistical significance; for **(H)**, asterisks color-coded as result curves indicate statistical significance (infant *n* = 40, adult *n* = 20, two-tailed sign-permutation tests, *P* < .05, FDR-corrected). For **(H)**, the black asterisks indicate significant difference between age groups (infant *n* = 40, adult *n*= 20, two-tailed Mann-Whitney *U* tests, *P* < .05, FDR-corrected).

**Figure S4.**
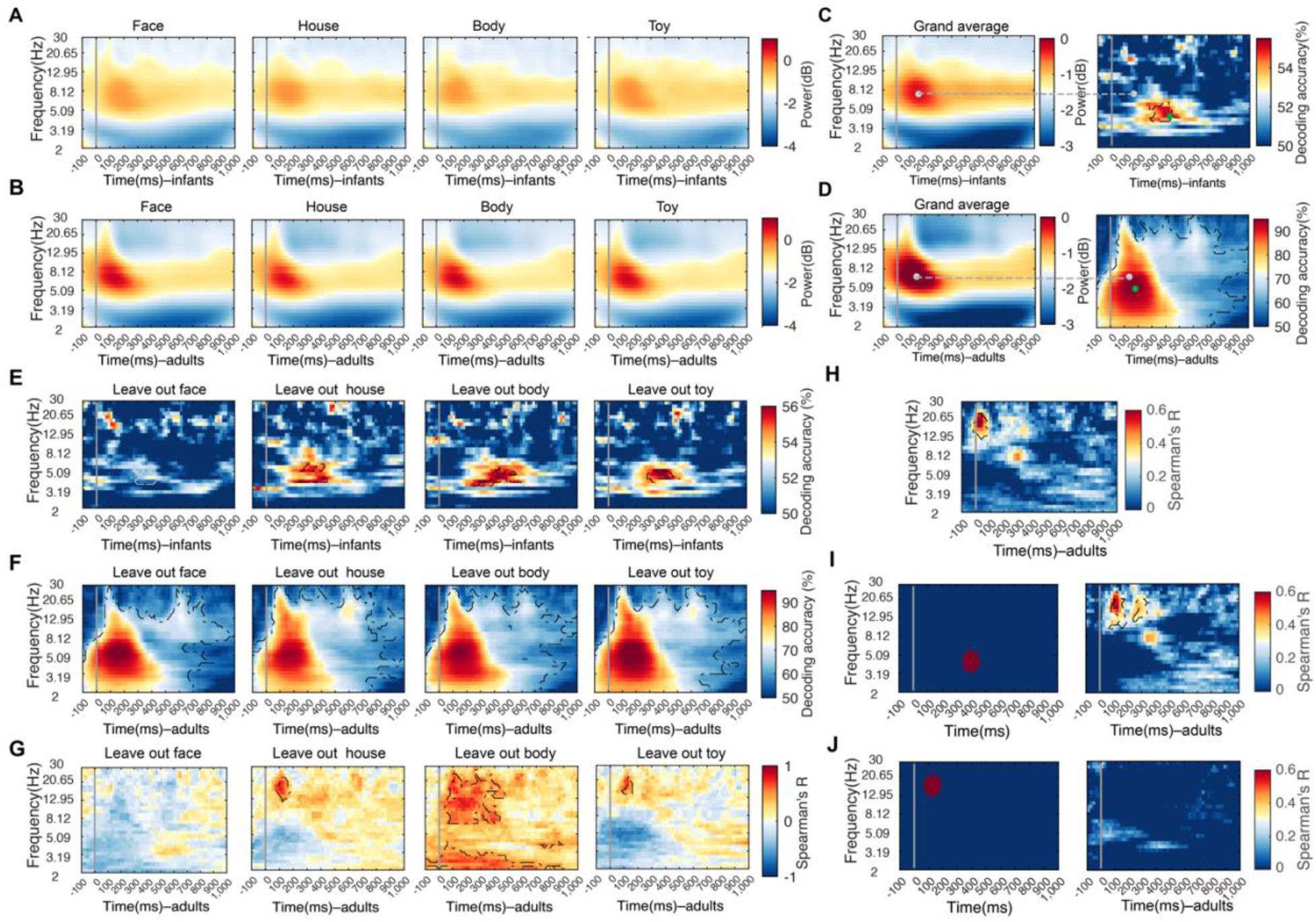
Analysis of Spectral Characterization of Infant and Adult Category Representations, related to Figure 4. **(A,B)** Category-specific EEG signal power resolved in frequency and time for infants **(A)** and for adults **(B)**. We decomposed the EEG time series using complex Morlet wavelets and expressed the spectrum of the power change versus baseline (from −300 to −100ms) during image presentation in decibel (dB). The dB power values was averaged across EEG channels for infants **(A)** and adults **(B)** as a function of time (from −100 to 1,000ms, in 20ms-steps) and frequency (from 2 to 30Hz, logarithmically spaced in 30 bins). **(C,D)** Visualization of the relationship between peaks in the grand-averaged EEG signal power and in the classification, infants in **(C)** and adults in **(D)**. We visualized the relationship between peaks (indicated by gray dots) in power in the time-frequency resolved signals (left panels) and peaks (indicated by green dots) in classification accuracy for category classification based on such signals (right panels in **C,D**). We observed a dissociation in frequency and timing. **(E,F)** Robustness of results in time- and frequency-resolved MVPA when leaving out each one of the four categories. The results in **(E,F)** are corresponding to the result in **Figure 4B,C**. For infants **(E)**, we observe specific clusters in the theta band with similar peak latency and frequency as in the main analysis, except in the leave-face-out analysis. For adults **(F)**, we observe a cluster extended across the whole frequency range and time course investigated as for the main analysis. **(G)** Robustness of results using RSA to link oscillation-based visual category representations in infants and adults. The results in **(G)** are equivalent to the analysis in **Figure 4E**, except that RDMs reduced by one column and row (corresponding to the category left out) were used, and infant oscillatory RDMs were created by averaging decoding accuracy matrices based on the extent of the cluster in the infant data, for each leave-category out analysis separately. The analyses yielded significant clusters of diverse shape and extent but in each case overlapping with the cluster in the alpha/beta range of the main analysis. For the analysis leaving faces out (the left most column) no such cluster existed, prohibiting this approach. In this case, we used the cluster extent from the main analyses instead as a stand-in. **(H–J)** Control analysis for using RSA to link oscillation-based visual category representations in infants and adults. The results in **(H–J)** are corresponding to the result in **Figure 4E**. **(H)** We linked oscillation-based object category representations in infants and adults as in the main analysis with the only differences that we aggregated data for the infant oscillatory RDMs based on the 95% confidence interval around peak frequency rather than cluster extent. Note that the 95% confidence interval is broad and thus this analysis is less specific in frequency than the main analysis. Nevertheless, we found a comparable pattern of results, with a main cluster of significant correlations around a peak at 17.13Hz (11.79–20.65Hz) and at 120ms (120–300ms). Detailed statistical information is listed in **Table S4C**. **(I,J)** We aggregated infants’ RDMs from time- and frequency-resolved classification within a radius of three time-points and frequency step combinations around the theta peak (**I**, left) and beta peak (**J**, left). RDMs from the infant theta peak were found to be similar to representations in adults in the alpha/beta range (left panel), reproducing the main analysis. RDMs from the infant alpha/beta peak were not significantly related to adult representations. Notes: The gray vertical lines indicate onset of image presentation. Black outlines indicate time- and frequency-point combinations with significant results (infants *n* = 40, adults *n* = 20, right-tailed permutation test, cluster-defining threshold *P* < .005, corrected significance level *P* < .05).

**Table S1.**
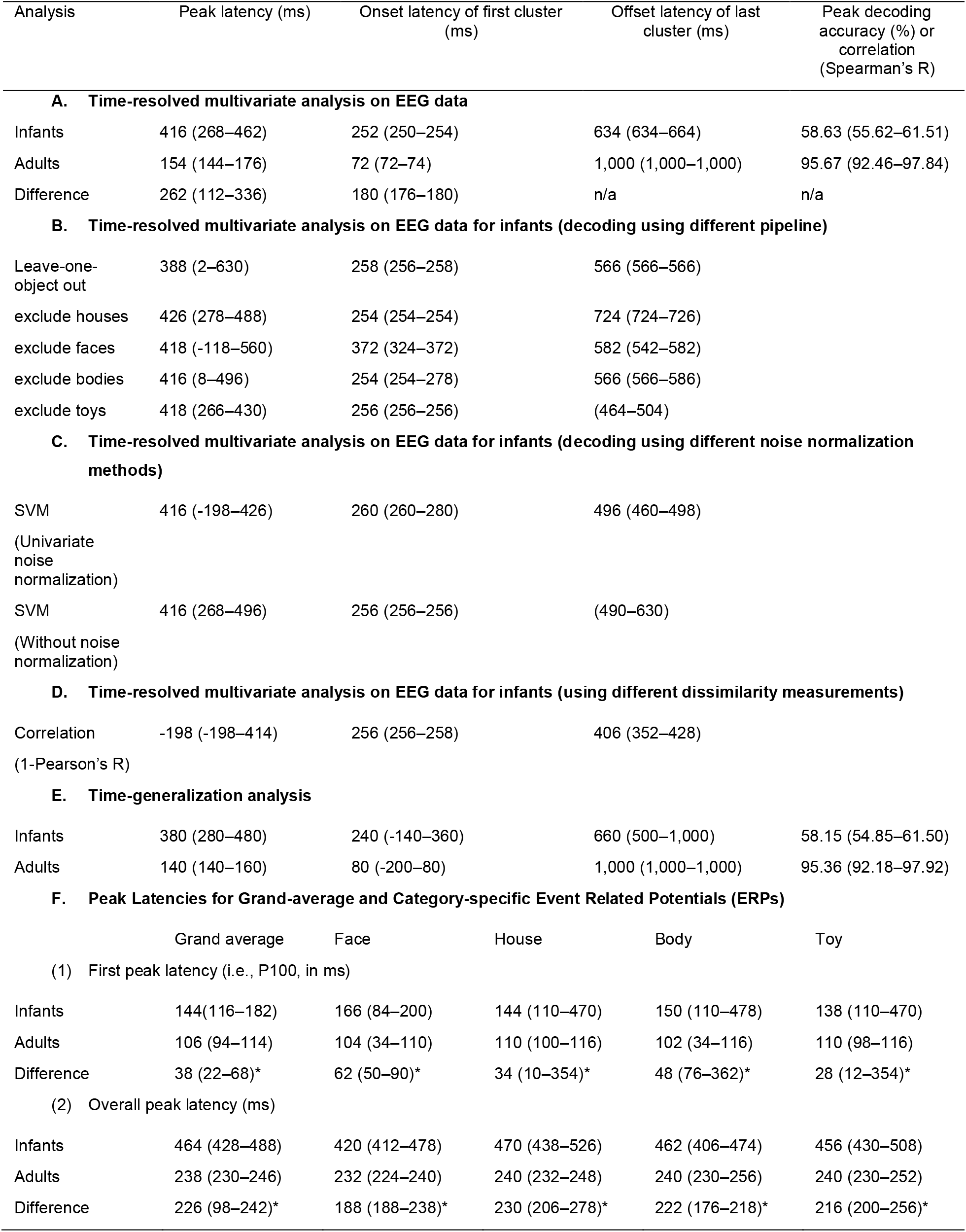
Statistical Details of the Time-resolved Multivariate Pattern EEG Classification and Event Related Potentials (ERPs), related to Figure 1 and Figure S1. The table enumerates the conducted analysis in the 1^st^ column (from **A** to **F**). For (**A–E**), the table lists the peak latency within a significant cluster (2^nd^ column), the onset latency of the first significant cluster (3^rd^ column), the offset of the last significant cluster (4^th^ column), and decoding accuracy (5^th^ column). For (**F**), the table lists **(1)** the first positive peak (i.e., P100) latency and **(2)** peak-latencies for event related potentials based on O1, O2 P7 and P8 electrodes. Results are given for the grand-average (1st column) and each category separately (2nd to 5th column). Notes: All values are averages across subjects (infant *n* = 40, adult *n* = 20), with 95% confidence intervals are reported in brackets. In (**F**), the stars after difference values indicate statistically significant difference (i.e., no overlap of 95% CIs between the two groups).

**Table S2.**
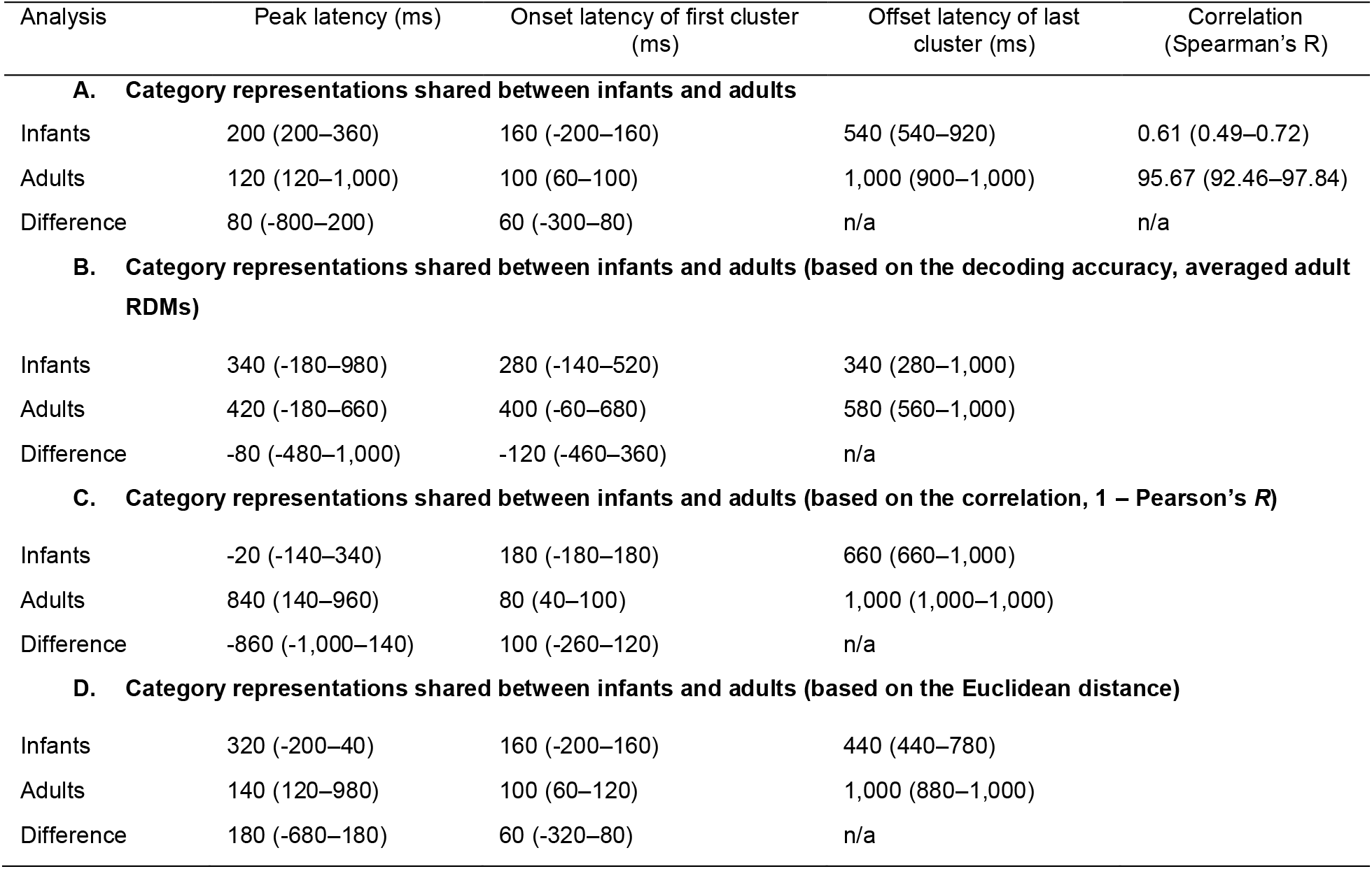
Statistical Details of the Across-Population Representational Similarity Analysis, related to Figure 2 and Figure S2. The table enumerates the conducted analysis in the 1^st^ column (from **A** to **D**), the peak latency within a significant cluster (2^nd^ column), the onset latency of the first significant cluster (3^rd^ column), the offset of the last significant cluster (4^th^ column), and correlation at peak (5^th^ column). Notes: All values are averages across subjects (infant *n* = 40, adult *n* = 20), with 95% confidence intervals are reported in brackets.

**Table S3.**
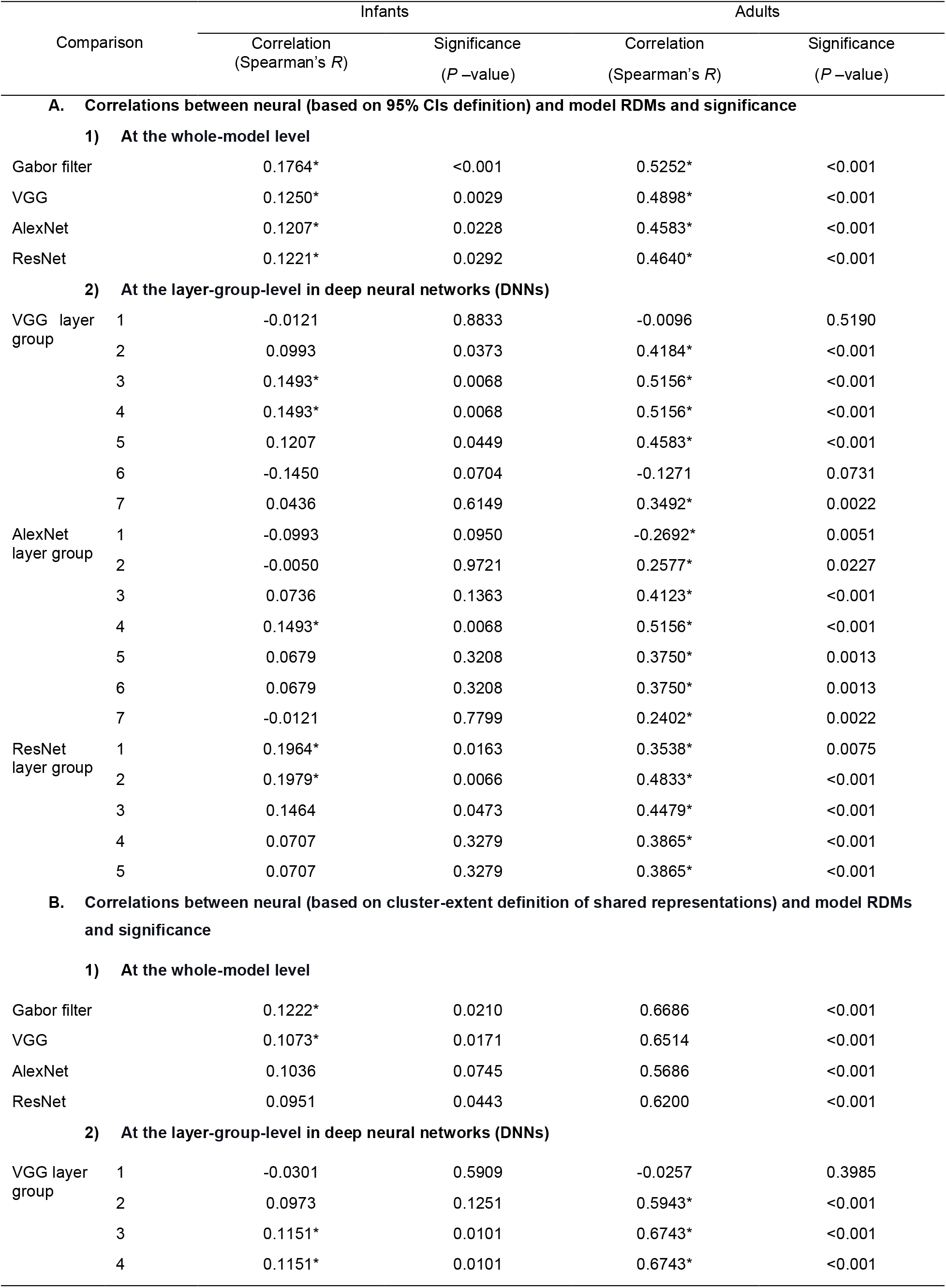
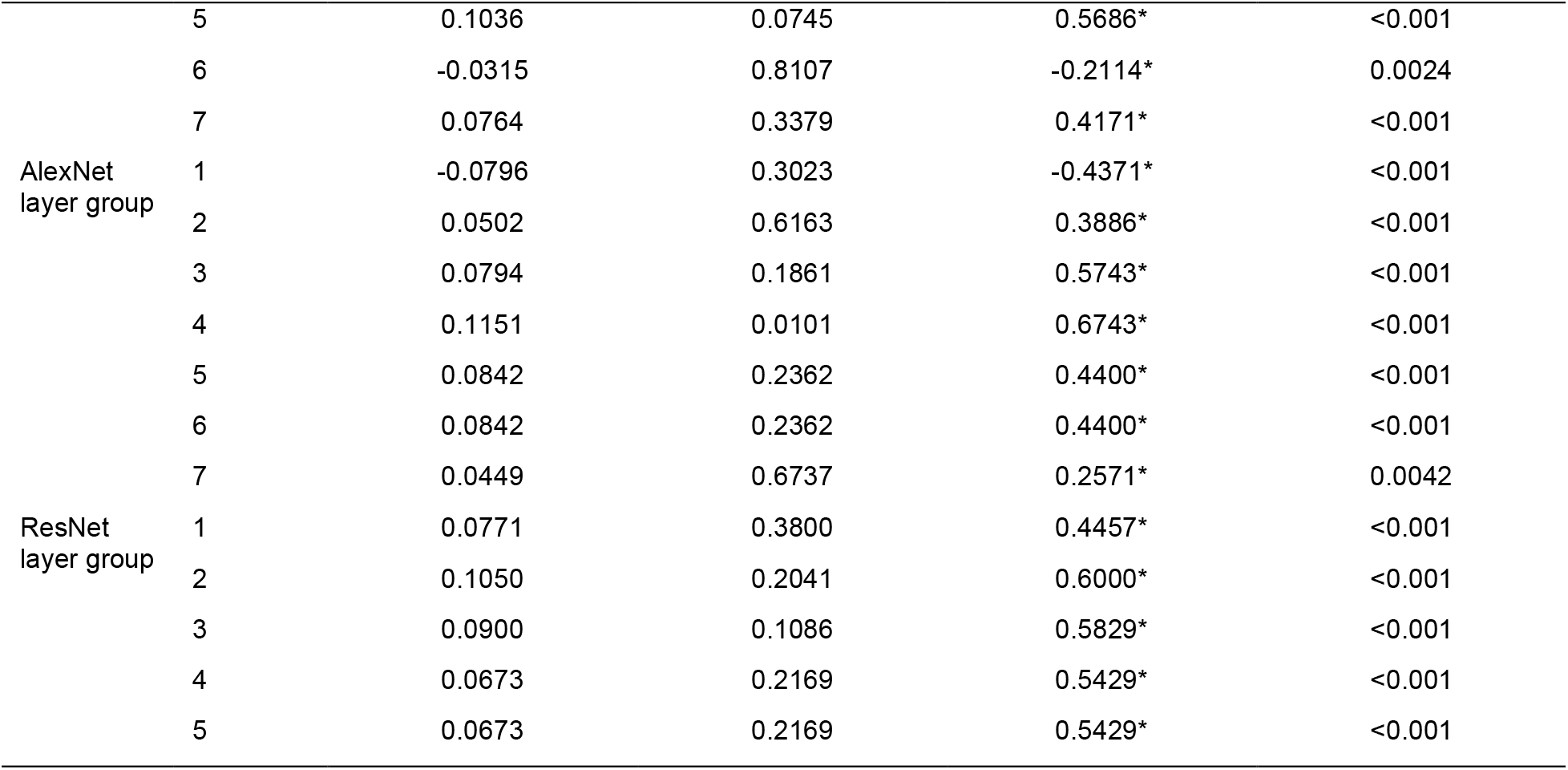
Correlations between Neural and Model RDMs and Significance, related to Figure 3 and Figure S3. The table reports the correlation between model and neural RDMs (defined by temporal extent of 95% CIs around the peak) and computational model RDMs at the **(A)** whole model and **(B)** layer-group level and its significance (infant *n* = 40, adult *n* = 20, two-tailed sign permutation tests, *P* < .05, FDR-corrected, significant correlations are indicated by a star).

**Table S4.**
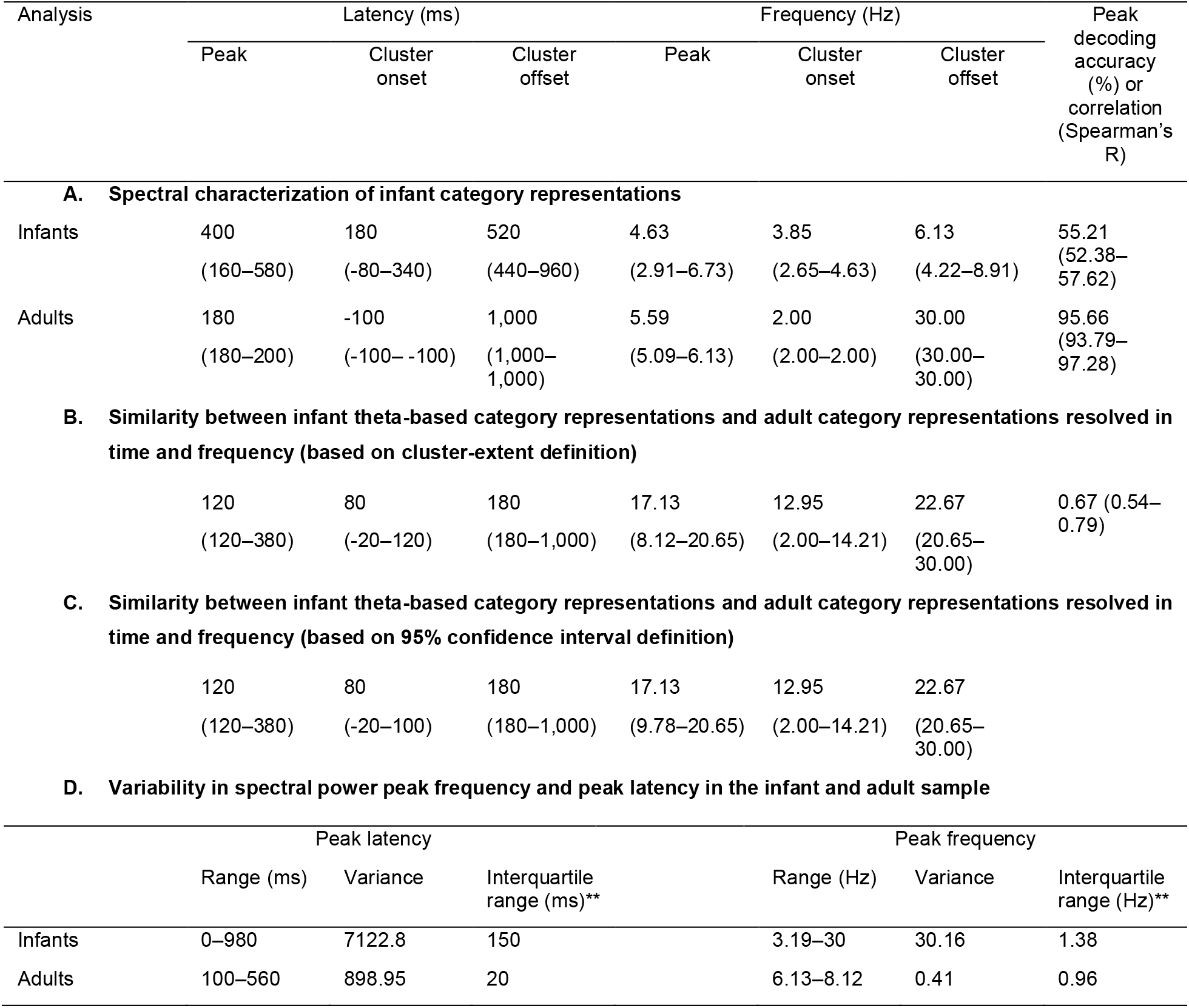
Statistical Details of Spectral Analyses and the Variability in Spectral Power, related to Figure 4 and Figure S4. The table enumerates the statistical details for spectral analysis and the variability in spectral power for infants and adults. The conducted analysis was shown in 1^st^ column. For **(A–C)**, we listed the peak latency within a significant cluster (2^nd^ column), the onset latency of the first significant cluster (3^rd^ column), and the offset of the last significant cluster (4^th^ column), the peak frequency within a significant cluster (5th column), the onset frequency of the first significant cluster (6th column), the offset frequency of the last significant cluster (7th column), and decoding accuracy or correlation at peak (8th column), 95% confidence intervals are reported in brackets. All values are averages across subjects (infant n = 40, adult n = 20), with 95% confidence intervals in brackets. For **(D)**, we listed the range, variance and interquartile range of the peak frequency and peak latency in the spectral power for both infants and adults. Consistently across all the three listed measures, for both time and frequency dimensions, adults show lower variability than infants. **Interquartile range is the difference between the 75th and 25th percentiles.

## REFERENCES

1. Potter, M. (1975). Meaning in visual search. Science 187, 965–966. 10.1126/science.1145183.

2. Thorpe, S., Fize, D., and Marlot, C. (1996). Speed of processing in the human visual system. Nature 381, 520–522. 10.1038/381520a0.

3. de Haan, M., and Nelson, C.A. (1999). Brain activity differentiates face and object processing in 6-month-old infants. Developmental Psychology 35, 1113–1121. 10.1037/0012-1649.35.4.1113.

4. Maurer, D., Lewis, T.L., Brent, H.P., and Levin, A.V. (1999). Rapid Improvement in the Acuity of Infants After Visual Input. Science 286, 108–110. 10.1126/science.286.5437.108.

5. Mareschal, D., and Quinn, P.C. (2001). Categorization in infancy. Trends in Cognitive Sciences 5, 443–450. 10.1016/S1364-6613(00)01752-6.

6. Pascalis, O., Haan, M. de, and Nelson, C.A. (2002). Is Face Processing Species-Specific During the First Year of Life? Science 296, 1321–1323. 10.1126/science.1070223.

7. Aslin, R.N. (2007). What’s in a look? Dev Sci 10, 48–53. 10.1111/J.1467-7687.2007.00563.X.

8. Aslin, R.N., and Fiser, J. (2005). Methodological challenges for understanding cognitive development in infants. Trends in Cognitive Sciences 9, 92–98. 10.1016/j.tics.2005.01.003.

9. Cichy, R.M., Pantazis, D., and Oliva, A. (2014). Resolving human object recognition in space and time. Nat Neurosci 17, 455–462. 10.1038/nn.3635.

10. Haxby, J.V., Gobbini, M.I., Furey, M.L., Ishai, A., Schouten, J.L., and Pietrini, P. (2001). Distributed and overlapping representations of faces and objects in ventral temporal cortex. Science 293, 2425–2430.

11. Grill-Spector, K., and Malach, R. (2004). The Human Visual Cortex. Annual Review of Neuroscience 27, 649–677. 10.1146/annurev.neuro.27.070203.144220.

12. Yamins, D.L.K., and DiCarlo, J.J. (2016). Using goal-driven deep learning models to understand sensory cortex. Nature Neuroscience 19, 356–365. 10.1038/nn.4244.

13. Singer, W. (1993). Synchronization of Cortical Activity and its Putative Role in Information Processing and Learning. Annual Review of Physiology 55, 349–374. 10.1146/annurev.ph.55.030193.002025.

14. Fries, P. (2015). Rhythms for Cognition: Communication through Coherence. Neuron 88, 220–235. 10.1016/j.neuron.2015.09.034.

15. Spriet, C., Abassi, E., Hochmann, J.-R., and Papeo, L. (2022). Visual object categorization in infancy. PNAS 119. 10.1073/pnas.2105866119.

16. Rakison, D.H., and Poulin-Dubois, D. (2001). Developmental origin of the animate–inanimate distinction. Psychological Bulletin 127, 209–228. 10.1037/0033-2909.127.2.209.

17. Stahl, A.E., and Feigenson, L. (2015). Observing the unexpected enhances infants’ learning and exploration. Science 348, 91–94. 10.1126/science.aaa3799.

18. Martin, A., Wiggs, C.L., Ungerleider, L.G., and Haxby, J.V. (1996). Neural correlates of category-specific knowledge. Nature 379, 649–652.

19. Deen, B., Richardson, H., Dilks, D.D., Takahashi, A., Keil, B., Wald, L.L., Kanwisher, N., and Saxe, R. (2017). Organization of high-level visual cortex in human infants. Nature Communications 8, 13995. 10.1038/ncomms13995.

20. Kosakowski, H.L., Cohen, M.A., Takahashi, A., Keil, B., Kanwisher, N., and Saxe, R. (2022). Selective responses to faces, scenes, and bodies in the ventral visual pathway of infants. Current Biology 32, 265–274.e5. 10.1016/j.cub.2021.10.064.

21. Arcaro, M.J., and Livingstone, M.S. (2017). A hierarchical, retinotopic proto-organization of the primate visual system at birth. eLife 6, e26196. 10.7554/eLife.26196.

22. Arcaro, M.J., Schade, P.F., and Livingstone, M.S. (2019). Body map proto-organization in newborn macaques. PNAS 116, 24861–24871. 10.1073/pnas.1912636116.

23. Livingstone, M.S., Vincent, J.L., Arcaro, M.J., Srihasam, K., Schade, P.F., and Savage, T. (2017). Development of the macaque face-patch system. Nature Communications 8, 14897. 10.1038/ncomms14897.

24. Ellis, C.T., Skalaban, L.J., Yates, T.S., Bejjanki, V.R., Córdova, N.I., and Turk-Browne, N.B. (2020). Re-imagining fMRI for awake behaving infants. Nature Communications 11, 4523. 10.1038/s41467-020-18286-y.

25. Yates, T.S., Ellis, C.T., and Turk-Browne, N.B. (2021). The promise of awake behaving infant fMRI as a deep measure of cognition. Current Opinion in Behavioral Sciences 40, 5–11. 10.1016/j.cobeha.2020.11.007.

26. Hoehl, S. (2016). The development of category specificity in infancy – What can we learn from electrophysiology? Neuropsychologia 83, 114–122. 10.1016/j.neuropsychologia.2015.08.021.

27. Conte, S., Richards, J.E., Guy, M.W., Xie, W., and Roberts, J.E. (2020). Face-sensitive brain responses in the first year of life. NeuroImage 211, 116602. 10.1016/j.neuroimage.2020.116602.

28. Kriegeskorte, N. (2008). Representational similarity analysis – connecting the branches of systems neuroscience. Front. Sys. Neurosci. 2, 4. doi: 10.3389/neuro.06.004.2008.

29. Grossmann, T., Gliga, T., Johnson, M.H., and Mareschal, D. (2009). The neural basis of perceptual category learning in human infants. J Cogn Neurosci 21, 2276–2286. 10.1162/jocn.2009.21188.

30. Quinn, P.C., Westerlund, A., and Nelson, C.A. (2006). Neural Markers of Categorization in 6-Month-Old Infants. Psychol Sci 17, 59–66. 10.1111/j.1467-9280.2005.01665.x.

31. Marinović, V., Hoehl, S., and Pauen, S. (2014). Neural correlates of human–animal distinction: An ERP-study on early categorical differentiation with 4-and 7-month-old infants and adults. Neuropsychologia 60, 60–76. 10.1016/j.neuropsychologia.2014.05.013.

32. Lee, J., Birtles, D., Wattam-Bell, J., Atkinson, J., and Braddick, O. (2012). Latency Measures of Pattern-Reversal VEP in Adults and Infants: Different Information from Transient P1 Response and Steady-State Phase. Investigative Ophthalmology & Visual Science 53, 1306–1314. 10.1167/iovs.11-7631.

33. McCulloch, D.L., and Skarf, B. (1991). Development of the human visual system: monocular and binocular pattern VEP latency. Investigative Ophthalmology & Visual Science 32, 2372–2381.

34. Moskowitz, A., and Sokol, S. (1983). Developmental changes in the human visual system as reflected by the latency of the pattern reversal VEP. Electroencephalogr Clin Neurophysiol 56, 1–15. 10.1016/0013-4694(83)90002-0.

35. Haynes, J.-D. (2015). A Primer on Pattern-Based Approaches to fMRI: Principles, Pitfalls, and Perspectives. Neuron 87, 257–270. 10.1016/j.neuron.2015.05.025.

36. Grootswagers, T., Wardle, S.G., and Carlson, T.A. (2016). Decoding Dynamic Brain Patterns from Evoked Responses: A Tutorial on Multivariate Pattern Analysis Applied to Time Series Neuroimaging Data. Journal of Cognitive Neuroscience 29, 677–697. 10.1162/jocn_a_01068.

37. Carlson, T., Tovar, D.A., Alink, A., and Kriegeskorte, N. (2013). Representational dynamics of object vision: The first 1000 ms. Journal of Vision 13, 1–19. 10.1167/13.10.1.

38. Ritchie, J.B., Tovar, D.A., and Carlson, T.A. (2015). Emerging Object Representations in the Visual System Predict Reaction Times for Categorization. PLoS Comput Biol 11, e1004316. 10.1371/journal.pcbi.1004316.

39. King, J.-R., and Dehaene, S. (2014). Characterizing the dynamics of mental representations: the temporal generalization method. Trends in Cognitive Sciences 18, 203–210. 10.1016/j.tics.2014.01.002.

40. Deoni, S.C.L., Mercure, E., Blasi, A., Gasston, D., Thomson, A., Johnson, M., Williams, S.C.R., and Murphy, D.G.M. (2011). Mapping Infant Brain Myelination with Magnetic Resonance Imaging. J. Neurosci. 31, 784–791. 10.1523/JNEUROSCI.2106-10.2011.

41. Huttenlocher, P.R., and Dabholkar, A.S. (1997). Regional differences in synaptogenesis in human cerebral cortex. Journal of Comparative Neurology 387, 167–178. https://doi.org/10.1002/(SICI)1096-9861(19971020)387:2<167::AID-CNE1>3.0.CO;2-Z.

42. Johnson, M.H. (2001). Functional brain development in humans. Nature Reviews Neuroscience 2, 475–483. 10.1038/35081509.

43. Kuhl, P.K., Williams, K.A., Lacerda, F., Stevens, K.N., and Lindblom, B. (1992). Linguistic experience alters phonetic perception in infants by 6 months of age. Science 255, 606–608. 10.1126/science.1736364.

44. Spelke, E.S., and Kinzler, K.D. (2007). Core knowledge. Developmental Science 10, 89–96. https://doi.org/10.1111/j.1467-7687.2007.00569.x.

45. Cichy, R.M., and Oliva, A. (2020). A M/EEG-fMRI Fusion Primer: Resolving Human Brain Responses in Space and Time. Neuron 107, 772–781. 10.1016/j.neuron.2020.07.001.

46. Cichy, R.M., Khosla, A., Pantazis, D., Torralba, A., and Oliva, A. (2016). Comparison of deep neural networks to spatio-temporal cortical dynamics of human visual object recognition reveals hierarchical correspondence. Scientific Reports 6, 27755. 10.1038/srep27755.

47. Jones, J.P., and Palmer, L.A. (1987). An evaluation of the two-dimensional Gabor filter model of simple receptive fields in cat striate cortex. Journal of Neurophysiology 58, 1233–1258. 10.1152/jn.1987.58.6.1233.

48. Simonyan, K., and Zisserman, A. (2015). Very Deep Convolutional Networks for Large-Scale Image Recognition. arXiv:1409.1556 [cs].

49. Kietzmann, T.C., McClure, P., and Kriegeskorte, N. (2019). Deep Neural Networks in Computational Neuroscience. Oxford Research Encyclopedia of Neuroscience. 10.1093/acrefore/9780190264086.013.46.

50. Graumann, M., Ciuffi, C., Dwivedi, K., Roig, G., and Cichy, R.M. (2022). The spatiotemporal neural dynamics of object location representations in the human brain. Nat Hum Behav 6, 796–811. 10.1038/s41562-022-01302-0.

51. Andrews, T.J., Clarke, A., Pell, P., and Hartley, T. (2010). Selectivity for low-level features of objects in the human ventral stream. NeuroImage 49, 703–711. 10.1016/j.neuroimage.2009.08.046.

52. Nasr, S., Echavarria, C.E., and Tootell, R.B.H. (2014). Thinking Outside the Box: Rectilinear Shapes Selectively Activate Scene-Selective Cortex. J. Neurosci. 34, 6721–6735. 10.1523/JNEUROSCI.4802-13.2014.

53. Oliva, A., and Torralba, A. (2006). Chapter 2 Building the gist of a scene: the role of global image features in recognition. In Visual Perception - Fundamentals of Awareness: Multi- Sensory Integration and High-Order Perception (Elsevier), pp. 23–36. 10.1016/S0079-6123(06)55002-2.

54. Kiorpes, L. (2016). The Puzzle of Visual Development: Behavior and Neural Limits. J. Neurosci. 36, 11384–11393. 10.1523/JNEUROSCI.2937-16.2016.

55. Peterzell, D.H., Werner, J.S., and Kaplan, P.S. (1995). Individual differences in contrast sensitivity functions: Longitudinal study of 4-, 6-and 8-month-old human infants. Vision Research 35, 961–979. 10.1016/0042-6989(94)00117-5.

56. Campbell, F.W., and Robson, J.G. (1968). Application of fourier analysis to the visibility of gratings. J Physiol 197, 551–566. 10.1113/jphysiol.1968.sp008574.

57. Kar, K., Kubilius, J., Schmidt, K., Issa, E.B., and DiCarlo, J.J. (2019). Evidence that recurrent circuits are critical to the ventral stream’s execution of core object recognition behavior. Nat Neurosci 22, 974–983. 10.1038/s41593-019-0392-5.

58. Kar, K., and DiCarlo, J.J. (2021). Fast recurrent processing via ventrolateral prefrontal cortex is needed by the primate ventral stream for robust core visual object recognition. Neuron 109, 164–176.

59. Tang, H., Schrimpf, M., Lotter, W., Moerman, C., Paredes, A., Caro, J.O., Hardesty, W., Cox, D., and Kreiman, G. (2018). Recurrent computations for visual pattern completion. PNAS 115, 8835–8840. 10.1073/pnas.1719397115.

60. Lupyan, G., Abdel Rahman, R., Boroditsky, L., and Clark, A. (2020). Effects of Language on Visual Perception. Trends in Cognitive Sciences 24, 930–944. 10.1016/j.tics.2020.08.005.

61. de Heering, A., and Rossion, B. (2015). Rapid categorization of natural face images in the infant right hemisphere. eLife 4, e06564. 10.7554/eLife.06564.

62. Hassabis, D., Kumaran, D., Summerfield, C., and Botvinick, M. (2017). Neuroscience-Inspired Artificial Intelligence. Neuron 95, 245–258. 10.1016/j.neuron.2017.06.011.

63. Berger, H. (1929). Über das Elektrenkephalogramm des Menschen. Archiv f. Psychiatrie 87, 527–570. 10.1007/BF01797193.

64. Buzsáki, G., and Draguhn, A. (2004). Neuronal Oscillations in Cortical Networks. Science 304, 1926–1929. 10.1126/science1099745.

65. Uhlhaas, P.J., Roux, F., Rodriguez, E., Rotarska-Jagiela, A., and Singer, W. (2010). Neural synchrony and the development of cortical networks. Trends in Cognitive Sciences 14, 72–80. 10.1016/j.tics.2009.12.002.

66. Marshall, P.J., Bar-Haim, Y., and Fox, N.A. (2002). Development of the EEG from 5 months to 4 years of age. Clinical Neurophysiology 113, 1199–1208. 10.1016/S1388-2457(02)00163-3.

67. Ward, L.M. (2003). Synchronous neural oscillations and cognitive processes. Trends in Cognitive Sciences 7, 553–559. 10.1016/j.tics.2003.10.012.

68. Buzsáki, G. (2005). Theta rhythm of navigation: link between path integration and landmark navigation, episodic and semantic memory. Hippocampus 15, 827–840. 10.1002/hipo.20113.

69. Jensen, O., and Mazaheri, A. (2010). Shaping Functional Architecture by Oscillatory Alpha Activity: Gating by Inhibition. Front. Hum. Neurosci. 4. 10.3389/fnhum.2010.00186.

70. Klimesch, W. (2012). Alpha-band oscillations, attention, and controlled access to stored information. Trends in Cognitive Sciences 16, 606–617. 10.1016/j.tics.2012.10.007.

71. Donoghue, T., Schaworonkow, N., and Voytek, B. (2022). Methodological considerations for studying neural oscillations. European Journal of Neuroscience 55, 3502–3527. 10.1111/ejn.15361.

72. Vidaurre, D., Cichy, R.M., and Woolrich, M.W. (2021). Dissociable Components of Information Encoding in Human Perception. Cerebral Cortex 31, 5664–5675. 10.1093/cercor/bhab189.

73. Tallon-Baudry, C., and Bertrand, O. (1999). Oscillatory gamma activity in humans and its role in object representation. Trends in Cognitive Sciences 3, 151–162. 10.1016/S1364-6613(99)01299-1.

74. Sauseng, P., Klimesch, W., Gruber, W.R., Hanslmayr, S., Freunberger, R., and Doppelmayr, M. (2007). Are event-related potential components generated by phase resetting of brain oscillations? A critical discussion. Neuroscience 146, 1435–1444. 10.1016/j.neuroscience.2007.03.014.

75. de Haan, M., Johnson, M.H., and Halit, H. (2003). Development of face-sensitive event-related potentials during infancy: a review. International Journal of Psychophysiology 51, 45–58. 10.1016/S0167-8760(03)00152-1.

76. Halit, H., Csibra, G., Volein, Á., and Johnson, M.H. (2004). Face-sensitive cortical processing in early infancy. Journal of Child Psychology and Psychiatry 45, 1228–1234. 10.1111/j.1469-7610.2004.00321.x.

77. Halit, H., de Haan, M., and Johnson, M.H. (2003). Cortical specialisation for face-processing: face sensitive event-related potential components in 3-and 12-month-old infants. NeuroImage 19, 1180–1193. 10.1016/S1053-8119(03)00076-4.

78. Hoehl, S., and Peykarjou, S. (2012). The early development of face processing — What makes faces special? Neurosci. Bull. 28, 765–788. 10.1007/s12264-012-1280-0.

79. Downing, P.E., Jiang, Y., Shuman, M., and Kanwisher, N. (2001). A Cortical Area Selective for Visual Processing of the Human Body. Science 293, 2470–2473. 10.1126/science.1063414.

80. Peelen, M.V., and Downing, P.E. (2005). Selectivity for the Human Body in the Fusiform Gyrus. J Neurophysiol 93, 603–608. 10.1152/jn.00513.2004.

81. Peykarjou, S., Pauen, S., and Hoehl, S. (2014). How do 9-month-old infants categorize human and ape faces? A rapid repetition ERP study. Psychophysiology 51, 866–878. 10.1111/psyp.12238.

82. Gliga, T., and Dehaene-Lambertz, G. (2007). Development of a view-invariant representation of the human head. Cognition 102, 261–288. 10.1016/j.cognition.2006.01.004.

83. Grill-Spector, K., and Weiner, K.S. (2014). The functional architecture of the ventral temporal cortex and its role in categorization. Nat Rev Neurosci 15, 536–548. 10.1038/nrn3747.

84. Oostenveld, R., Fries, P., Maris, E., and Schoffelen, J.-M. (2010). FieldTrip: Open Source Software for Advanced Analysis of MEG, EEG, and Invasive Electrophysiological Data. Computational Intelligence and Neuroscience 2011, e156869. https://doi.org/10.1155/2011/156869.

85. Tadel, F., Baillet, S., Mosher, J.C., Pantazis, D., and Leahy, R.M. (2011). Brainstorm: A User-Friendly Application for MEG/EEG Analysis. Computational Intelligence and Neuroscience 2011, 1–13. 10.1155/2011/879716.

86. Chang, C., and Lin, C. (2001). {LIBSVM}: a library for support vector machines.

87. Kriegeskorte, N., Goebel, R., and Bandettini, P. (2006). Information-based functional brain mapping. Proceedings of the National Academy of Sciences of the United States of America 103, 3863–3868. 10.1073/pnas.0600244103.

88. Kaiser, D., Oosterhof, N.N., and Peelen, M.V. (2016). The Neural Dynamics of Attentional Selection in Natural Scenes. J Neurosci 36, 10522–10528. 10.1523/JNEUROSCI.1385-16.2016.

89. Kriegeskorte, N., Mur, M., Ruff, D.A., Kiani, R., Bodurka, J., Esteky, H., Tanaka, K., and Bandettini, P.A. (2008). Matching Categorical Object Representations in Inferior Temporal Cortex of Man and Monkey. Neuron 60, 1126–1141. 10.1016/j.neuron.2008.10.043.

90. Kriegeskorte, N., and Kievit, R.A. (2013). Representational geometry: integrating cognition, computation, and the brain. Trends in Cognitive Sciences 17, 401–412. 10.1016/j.tics.2013.06.007.

91. Guggenmos, M., Sterzer, P., and Cichy, R.M. (2018). Multivariate pattern analysis for MEG: A comparison of dissimilarity measures. Neuroimage 173, 434–447. 10.1016/j.neuroimage.2018.02.044.

92. Kay, K.N., Naselaris, T., Prenger, R.J., and Gallant, J.L. (2008). Identifying natural images from human brain activity. Nature 452, 352–355. 10.1038/nature06713.

93. Simonyan, K., Vedaldi, A., and Zisserman, A. (2013). Deep Inside Convolutional Networks: Visualising Image Classification Models and Saliency Maps. arXiv:1312.6034 [cs].

94. Bau, D., Zhou, B., Khosla, A., Oliva, A., and Torralba, A. (2017). Network Dissection: Quantifying Interpretability of Deep Visual Representations. arXiv:1704.05796 [cs].

95. Güçlü, U., and Gerven, M.A.J. van (2015). Deep Neural Networks Reveal a Gradient in the Complexity of Neural Representations across the Ventral Stream. J. Neurosci. 35, 10005–10014. 10.1523/JNEUROSCI.5023-14.2015.

96. Eickenberg, M., Gramfort, A., Varoquaux, G., and Thirion, B. (2017). Seeing it all: Convolutional network layers map the function of the human visual system. NeuroImage 152, 184–194. 10.1016/j.neuroimage.2016.10.001.

97. Khaligh-Razavi, S.-M., and Kriegeskorte, N. (2014). Deep Supervised, but Not Unsupervised, Models May Explain IT Cortical Representation. PLoS Comput Biol 10, e1003915. 10.1371/journal.pcbi.1003915.

98. Yamins, D.L.K., Hong, H., Cadieu, C.F., Solomon, E.A., Seibert, D., and DiCarlo, J.J. (2014). Performance-optimized hierarchical models predict neural responses in higher visual cortex. PNAS 111, 8619–8624. 10.1073/pnas.1403112111.

99. Schrimpf, M., Kubilius, J., Lee, M.J., Ratan Murty, N.A., Ajemian, R., and DiCarlo, J.J. (2020). Integrative Benchmarking to Advance Neurally Mechanistic Models of Human Intelligence. Neuron 108, 413–423. 10.1016/j.neuron.2020.07.040.

100. Vedaldi, A., and Lenc, K. (2016). MatConvNet - Convolutional Neural Networks for MATLAB. arXiv:1412.4564 [cs].

101. DiCarlo, J.J., Zoccolan, D., and Rust, N.C. (2012). How Does the Brain Solve Visual Object Recognition? Neuron 73, 415–434. 10.1016/j.neuron.2012.01.010.

102. Nili, H., Wingfield, C., Walther, A., Su, L., Marslen-Wilson, W., and Kriegeskorte, N. (2014). A Toolbox for Representational Similarity Analysis. PLoS Comput Biol 10, e1003553. 10.1371/journal.pcbi.1003553.

103. Maris, E., and Oostenveld, R. (2007). Nonparametric statistical testing of EEG-and MEG-data. Journal of Neuroscience Methods 164, 177–190. 10.1016/j.jneumeth.2007.03.024.

104. Nichols, T.E., and Holmes, A.P. (2002). Nonparametric permutation tests for functional neuroimaging: A primer with examples. Hum. Brain Mapp. 15, 1–25. 10.1002/hbm.1058.

