## Supplemental information for "Visual category representations in the infant brain"

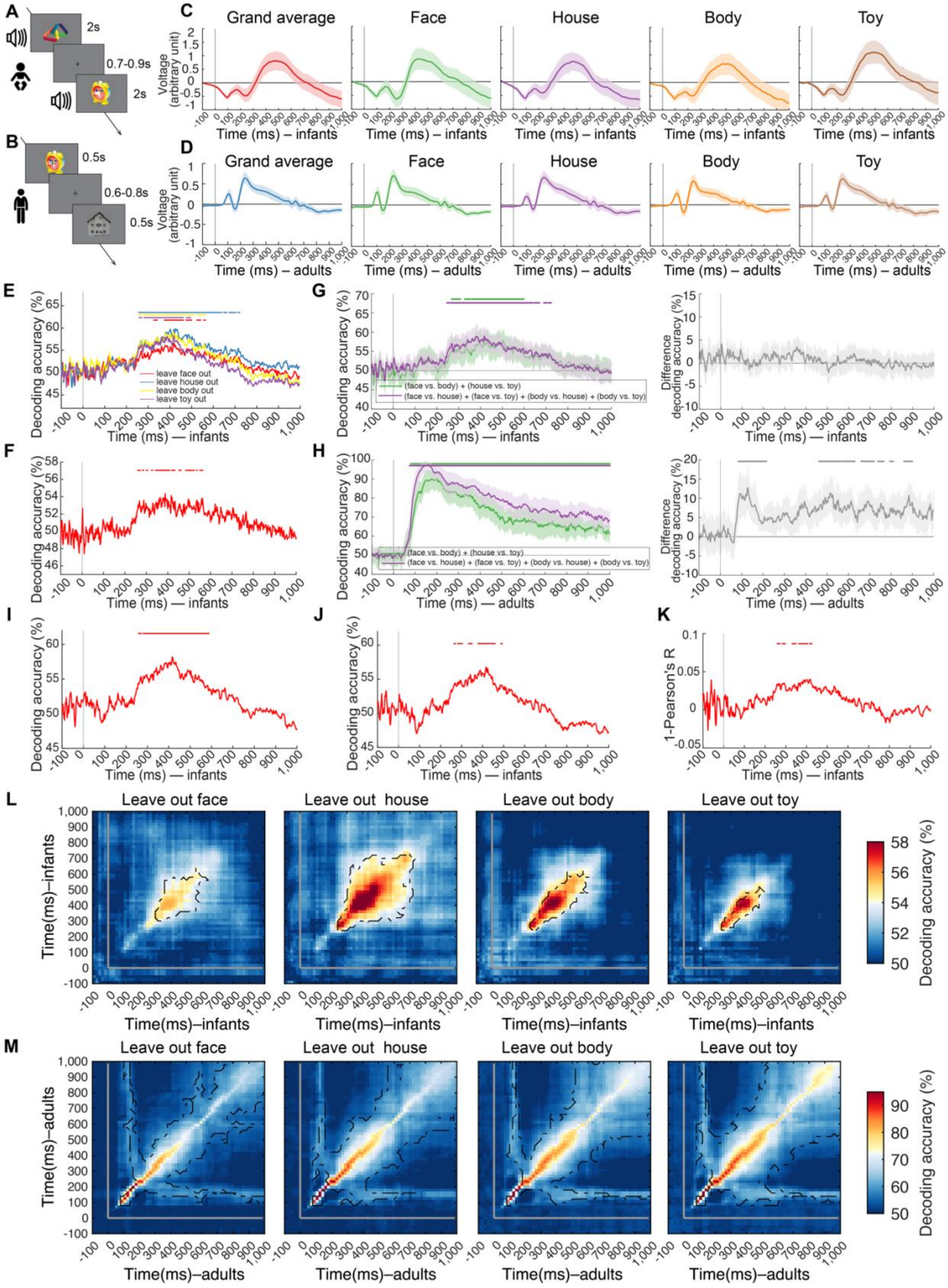
A
2s
0.7-0.9s
2s
B
0.5s
0.6-0.8s
0.5s
C
Grand average
Face
House
Body
Toy
Voltage (arbitrary unit)
Time (ms) – infants
D
Grand average
Face
House
Body
Toy
Voltage (arbitrary unit)
Time (ms) – adults
E
Decoding accuracy (%)
leave face out
leave house out
leave body out
leave toy out
Time (ms) — infants
F
Decoding accuracy (%)
Time (ms) — infants
G
Decoding accuracy (%)
(face vs. body) + (house vs. toy)
(face vs. house) + (face vs. toy) + (body vs. house) + (body vs. toy)
Time (ms) — infants
Difference decoding accuracy (%)
Time (ms) — infants
H
Decoding accuracy (%)
(face vs. body) + (house vs. toy)
(face vs. house) + (face vs. toy) + (body vs. house) + (body vs. toy)
Time (ms) — adults
Difference decoding accuracy (%)
Time (ms) — adults
I
Decoding accuracy (%)
Time (ms) — infants
J
Decoding accuracy (%)
Time (ms) — infants
K
1-Pearson's R
Time (ms) — infants
L
Leave out face
Leave out house
Leave out body
Leave out toy
Time(ms)–infants
Decoding accuracy (%)
M
Leave out face
Leave out house
Leave out body
Leave out toy
Time(ms)–adults
Decoding accuracy (%)

**Figure S1. Analysis of Category Representations in Time for Infants and Adults, related to Figure 1 and STAR Methods. (A,B)** Experimental paradigms. For the infant sample **(A)**, on each trial an object image was presented for 2s together with a randomly assigned 10 arbitrary sounds, followed by a variable inter-stimulus-interval (ISI) of 0.7s to 0.9s; for adult sample **(B)**, presentation parameters were adapted to increase stimulus repetitions and thus the signal-to-noise ratio. On each trial an object image was presented for 0.5s followed by a variable ISI of 0.6s to 0.8s. **(C,D)** Visualization of event related potentials (ERPs). We averaged baseline-normalized (mean removal and division by standard deviation of the baseline activity) trials (electrodes: O1, O2, P7, P8) across category (i.e., grand average) or by category (i.e., face, house, body and toy). The ERPs were plotted in **(C)** for infants and in **(D)** for adults. In each case, the reported ERPs were either the grand average across all categories or averaged for each category separately. P100 peak latencies were slightly but significantly delayed in infants (i.e., 22–68ms difference). Overall peak latency differences were moderate (i.e., 98–242ms difference). For detailed peak latencies and statistical information, see **Table S1F**. For visualization, results were smoothed within a 20-ms sliding window. Shaded areas above and below curves indicate 95% confidence intervals determined by bootstrapping the participant sample (1,000 iterations). The gray vertical line indicates the onset of image presentation. **(E,F)** Time-resolved multivariate analysis in infants do not depend on any particular object or category. To determine whether the pattern of results depended strongly on any one category, we averaged category classification results leaving out one of the four categories in turn. The results in **(E)** are equivalent to **Figure 1C**. To determine whether the time-resolved results depended strongly on any one object in the stimulus set, we performed category decoding using a leave-one-object-out cross-validation scheme. In detail, we trained the classifier using data from 31 of the 32 images of a category (e.g., 31 faces vs. 31 toys), and tested it on the left-out images (e.g., the 32nd face and toy). The results in **(F)** are equivalent to **Figure 1C**. Detailed statistical information for **(E,F)** is listed in **Table S1B**. **(G,H)** Results of time-resolved multivariate analysis held equally within and across the animacy distinction in both infants and adults. We grouped pairwise category classification results by the animacy of the categories classified. In detail, we averaged the results from 1) classifying within the animate (face vs. body) and within the inanimate (house vs. toy) division together (green curves); 2) classifying across the animate and the inanimate division (face vs. house, face vs. toy, body vs. house and body vs. toy) together (magenta curves). In infants **(G)**, we observed statistically significant results for classification within and across the animacy division separately (left panel) but the difference was not significant (right panel). In contrast and as expected (Carlson et al., 2013; Cichy et al., 2014); in adults **(H)**, we observed statistically significant results for classification both within and across the animacy division (left panel) and significantly higher results across the animacy division (right panel). It shows that infants represent finer distinctions than the basic animate vs. inanimate division and suggests that visual category representations do partly reflect the animacy division in adults, but not yet in our infant sample. The shaded margin on curves indicates 95% confidence intervals of decoding accuracy. **(I–K)** Results of time-resolved multivariate analysis in infants emerged equivalently for alternative analysis schemes. Category classification results did not strongly depend on particular preprocessing steps: results without **(I)** or with **(J)** univariate noise normalization (i.e. each trial and channel separately was z-transformed based on baseline activity from –200 to 0ms). Category differentiation showed an equivalent result pattern **(K)** when comparing category-specific EEG response patterns using 1-Pearson's R rather than decoding accuracy as a measure. Detailed statistical information for **(I–K)** is listed in **Table S1C,D**. **(L,M)** Results of time generalization analyses showed robustness in both infants and adults when leaving out each one of the four categories To determine whether the pattern of results depended strongly on any one category, we averaged category classification results leaving out one of the four categories in turn. The results in **(L,M)** are equivalent to **Figure 1G,H**, respectively. We observe a similar result pattern as in the main analysis in each case. Notes: The gray lines on plots indicate image onset. Rows of asterisks in **(E–K)** indicate significant time points. Black outlines in **(L,M)** indicate time point combinations with significant results (infant $n = 40$, adult $n = 20$, right-tailed sign permutation tests, cluster-defining threshold $P < .005$, corrected significance level $P < .05$).

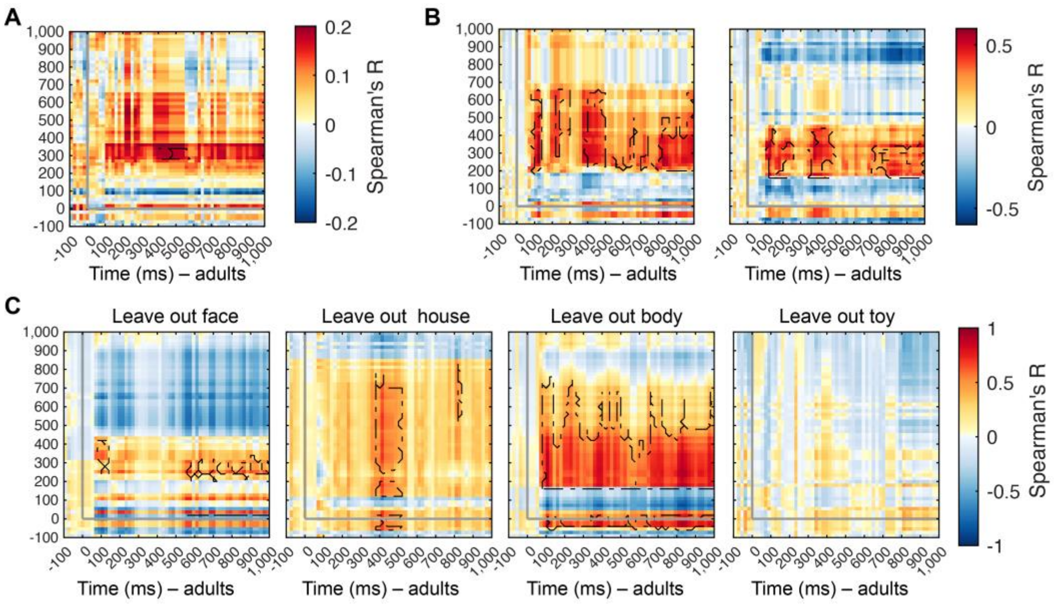


**Figure S2. Analysis of Shared Category Representations between Infants and Adults, related to Figure 2. (A,B)** Results of time-generalization RSA was similarly achieved for alternative processing and data aggregation choices. We compared RDMs (Spearman's *R*) in infants and adults averaging over adult participants, rather than infant participants and the result was shown in **(A)**. Detailed statistical information is listed in **Table S2B**. We used 1 – Pearson's *R* in (**B**, left) and Euclidean distance in (**B**, right) as a dissimilarity measure between brain patterns rather than decoding accuracy. Detailed statistical information is listed in **Table S2C,D**. **(C)** Results of time-generalization RSA did not depend on any particular category except on toys. To determine whether the pattern of results depended strongly on any one category, we left out one of the four categories in turn for RSA. We observe evidence for shared visual category representations in all analyses except the leave-toy-out analysis. Notes: The gray lines on plots indicate image onset. Black outlines indicate time point combinations with significant results (right-tailed sign permutation tests, cluster-defining threshold $P < .005$, corrected significance level $P < .05$).

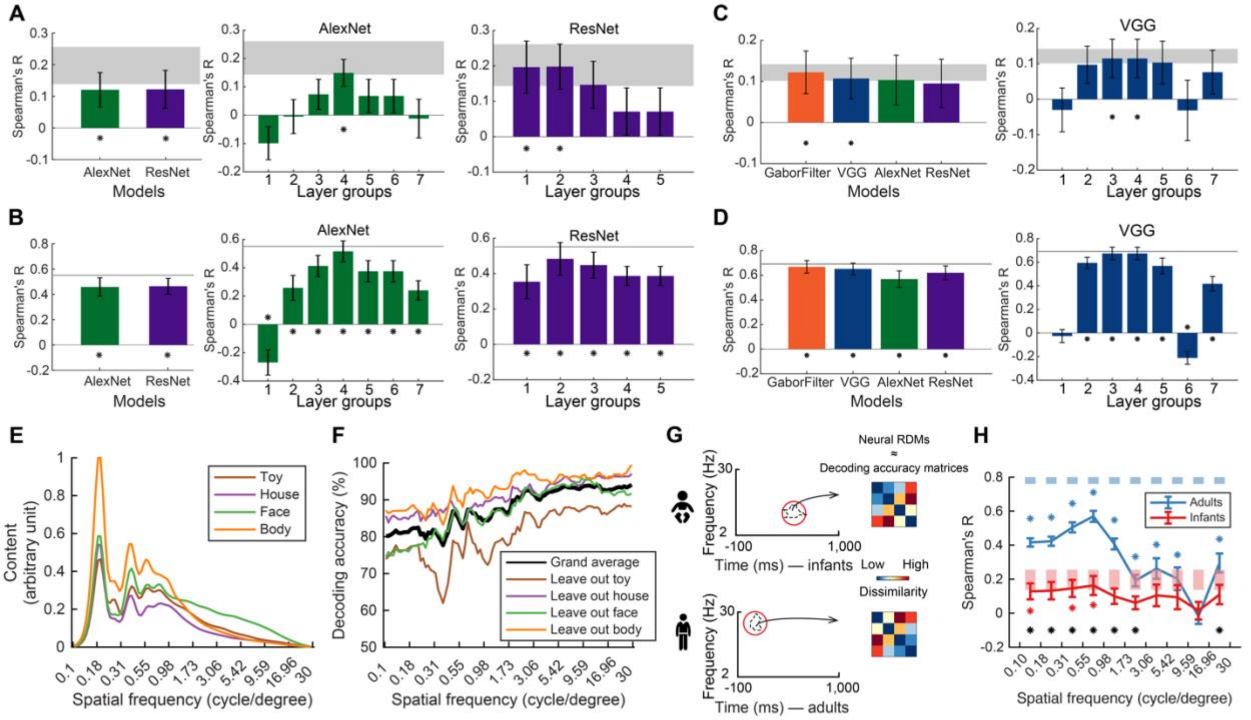


**Figure S3. Analysis of the Format of Category Representations in Infants and Adults, related to Figure 3. (A,B)** We assessed the format of category representations in infants and adults with relation to various DNNs (i.e., AlexNet and ResNet-50). Results for infants **(A)** and adults **(B)** at the whole-model level and DNN layer level. Statistical details (i.e., correlations and *P*-values) are in **Table S3A**. **(C,D)** We assessed the format of category representations using the significant shared representations, as shown in Figure 2B. For this, we defined neural RDMs as the average of decoding accuracy RDMs from time-resolved category classification in the temporal extent of the cluster indicating shared representations. Results for infants **(C)** and adults **(D)** at the whole-model level and at the DNN layer level for VGG models. Statistical details (i.e., correlations and P-values) are in **Table S3B**. **(E)** Spatial frequency spectrum in image categories. The panel showed the mean spatial frequency power in images for objects, ordered by category. Values are normalized such that the maximum value across categories is set to 1. The procedure was as follows. All images were filtered into one hundred bins logarithmically spaced between 0.1 and 30 cycles per degree visual angle using Butterworth band-pass filters. To compute image-specific spatial frequency power, each original image was first Fourier transformed into the frequency domain, multiplied by a 2-D frequency band-pass filter, and the power value was extracted as the spatial frequency content at that frequency bin. **(F)** Results of category classification from the spatial-frequency-specific power values of the images, averaged across all comparisons (grand average) and for each analysis leaving one category out. **(G)** Spatial-frequency-specific features in representations shared between adults and infants. For this, we extracted RDMs from the time-frequency classification analysis in the theta cluster in infants, and RDMs from the time-frequency classification analysis in adults at the location of the cluster indicating shared representations between infants and adults. We compared each in turn to spatial-frequency specific RDMs of the DNN. **(H)** Results indicate that adults represent features at all spatial frequencies, whereas infants do so only in low spatial frequencies. The conjunction of those results reveals that the features shared between infants and adults are in low spatial frequencies. Notes: For **(A–D,H)**, error bars represent standard errors of the mean. For **(A–D)**, asterisks indicate statistical significance; for **(H)**, asterisks color-coded as result curves indicate statistical significance (infant $n = 40$, adult $n = 20$, two-tailed sign-permutation tests, $P < .05$, FDR-corrected). For **(H)**, the black asterisks indicate significant difference between age groups (infant $n = 40$, adult $n = 20$, two-tailed Mann-Whitney $U$ tests, $P < .05$, FDR-corrected).

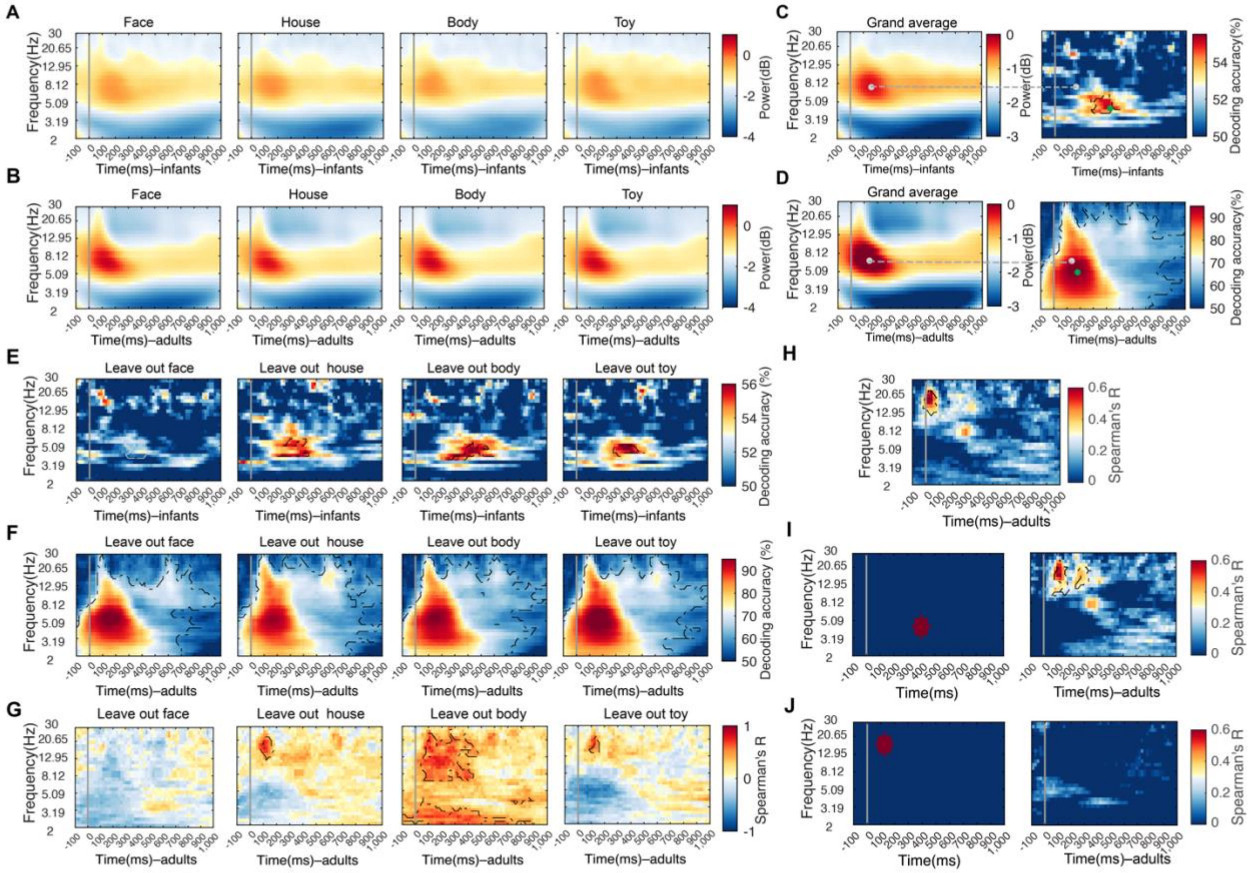


**Figure S4. Analysis of Spectral Characterization of Infant and Adult Category Representations, related to Figure 4. (A,B)** Category-specific EEG signal power resolved in frequency and time for infants **(A)** and for adults **(B)**. We decomposed the EEG time series using complex Morlet wavelets and expressed the spectrum of the power change versus baseline (from –300 to –100ms) during image presentation in decibel (dB). The dB power values was averaged across EEG channels for infants **(A)** and adults **(B)** as a function of time (from –100 to 1,000ms, in 20ms-steps) and frequency (from 2 to 30Hz, logarithmically spaced in 30 bins). **(C,D)** Visualization of the relationship between peaks in the grand-averaged EEG signal power and in the classification, infants in **(C)** and adults in **(D)**. We visualized the relationship between peaks (indicated by gray dots) in power in the time-frequency resolved signals (left panels) and peaks (indicated by green dots) in classification accuracy for category classification based on such signals (right panels in **C,D**). We observed a dissociation in frequency and timing. **(E,F)** Robustness of results in time- and frequency-resolved MVPA when leaving out each one of the four categories. The results in **(E,F)** are corresponding to the result in **Figure 4B,C**. For infants **(E)**, we observe specific clusters in the theta band with similar peak latency and frequency as in the main analysis, except in the leave-face-out analysis. For adults **(F)**, we observe a cluster extended across the whole frequency range and time course investigated as for the main analysis. **(G)** Robustness of results using RSA to link oscillation-based visual category representations in infants and adults. The results in **(G)** are equivalent to the analysis in **Figure 4E**, except that RDMs reduced by one column and row (corresponding to the category left out) were used, and infant oscillatory RDMs were created by averaging decoding accuracy matrices based on the extent of the cluster in the infant data, for each leave-category out analysis separately. The analyses yielded significant clusters of diverse shape and extent but in each case overlapping with the cluster in the alpha/beta range of the main analysis. For the analysis leaving faces out (the left most column) no such cluster existed, prohibiting this approach. In this case, we used the cluster extent from the main analyses instead as a stand-in. **(H–J)** Control analysis for using RSA to link oscillation-based visual category representations in infants and adults. The results in **(H–J)** are corresponding to the result in **Figure 4E. (H)** We linked oscillation-based object category representations in infants and adults as in the main analysis with the only differences that we aggregated data for the infant oscillatory RDMs based on the 95% confidence interval around peak frequency rather than cluster extent. Note that the 95% confidence interval is broad and thus this analysis is less specific in frequency than the main analysis. Nevertheless, we found a comparable pattern of results, with a main cluster of significant correlations around a peak at 17.13Hz (11.79–20.65Hz) and at 120ms (120–300ms). Detailed statistical information is listed in **Table S4C**. **(I,J)** We aggregated infants' RDMs from time- and frequency-resolved classification within a radius of three time -points and frequency step combinations around the theta peak **(I**, left**)** and beta peak **(J**, left**)**. RDMs from the infant theta peak were found to be similar to representations in adults in the alpha/beta range (left panel), reproducing the main analysis. RDMs from the infant alpha/beta peak were not significantly related to adult representations. Notes: The gray vertical lines indicate onset of image presentation. Black outlines indicate time- and frequency-point combinations with significant results (infants $n$ = 40, adults $n$ = 20, right-tailed permutation test, cluster-defining threshold $P$ < .005, corrected significance level $P$ < .05).

| Analysis | Peak latency (ms) | Onset latency of first cluster (ms) | Offset latency of last cluster (ms) | Peak decoding accuracy (%) or correlation (Spearman's R) |
|---|---|---|---|---|
| **A. Time-resolved multivariate analysis on EEG data** | | | | |
| Infants | 416 (268–462) | 252 (250–254) | 634 (634–664) | 58.63 (55.62–61.51) |
| Adults | 154 (144–176) | 72 (72–74) | 1,000 (1,000–1,000) | 95.67 (92.46–97.84) |
| Difference | 262 (112–336) | 180 (176–180) | n/a | n/a |
| **B. Time-resolved multivariate analysis on EEG data for infants (decoding using different pipeline)** | | | | |
| Leave-one-object out | 388 (2–630) | 258 (256–258) | 566 (566–566) | |
| exclude houses | 426 (278–488) | 254 (254–254) | 724 (724–726) | |
| exclude faces | 418 (-118–560) | 372 (324–372) | 582 (542–582) | |
| exclude bodies | 416 (8–496) | 254 (254–278) | 566 (566–586) | |
| exclude toys | 418 (266–430) | 256 (256–256) | (464–504) | |
| **C. Time-resolved multivariate analysis on EEG data for infants (decoding using different noise normalization methods)** | | | | |
| SVM (Univariate noise normalization) | 416 (-198–426) | 260 (260–280) | 496 (460–498) | |
| SVM (Without noise normalization) | 416 (268–496) | 256 (256–256) | (490–630) | |
| **D. Time-resolved multivariate analysis on EEG data for infants (using different dissimilarity measurements)** | | | | |
| Correlation (1-Pearson's R) | -198 (-198–414) | 256 (256–258) | 406 (352–428) | |
| **E. Time-generalization analysis** | | | | |
| Infants | 380 (280–480) | 240 (-140–360) | 660 (500–1,000) | 58.15 (54.85–61.50) |
| Adults | 140 (140–160) | 80 (-200–80) | 1,000 (1,000–1,000) | 95.36 (92.18–97.92) |

**F. Peak Latencies for Grand-average and Category-specific Event Related Potentials (ERPs)**

| | Grand average | Face | House | Body | Toy |
|---|---|---|---|---|---|
| (1) First peak latency (i.e., P100, in ms) | | | | | |
| Infants | 144(116–182) | 166 (84–200) | 144 (110–470) | 150 (110–478) | 138 (110–470) |
| Adults | 106 (94–114) | 104 (34–110) | 110 (100–116) | 102 (34–116) | 110 (98–116) |
| Difference | 38 (22–68)* | 62 (50–90)* | 34 (10–354)* | 48 (76–362)* | 28 (12–354)* |
| (2) Overall peak latency (ms) | | | | | |
| Infants | 464 (428–488) | 420 (412–478) | 470 (438–526) | 462 (406–474) | 456 (430–508) |
| Adults | 238 (230–246) | 232 (224–240) | 240 (232–248) | 240 (230–256) | 240 (230–252) |
| Difference | 226 (98–242)* | 188 (188–238)* | 230 (206–278)* | 222 (176–218)* | 216 (200–256)* |

**Table S1. Statistical Details of the Time-resolved Multivariate Pattern EEG Classification and Event Related Potentials (ERPs), related to Figure 1 and Figure S1.** The table enumerates the conducted analysis in the 1st column (from **A** to **F**). For (**A–E**), the table lists the peak latency within a significant cluster (2nd column), the onset latency of the first significant cluster (3rd column), the offset of the last significant cluster (4th column), and decoding accuracy

(5$^{th}$ column). For (**F**), the table lists **(1)** the first positive peak (i.e., P100) latency and **(2)** peak-latencies for event related potentials based on O1, O2 P7 and P8 electrodes. Results are given for the grand-average (1st column) and each category separately (2nd to 5th column). Notes: All values are averages across subjects (infant *n* = 40, adult *n* = 20), with 95% confidence intervals are reported in brackets. In (**F**), the stars after difference values indicate statistically significant difference (i.e., no overlap of 95% CIs between the two groups).

| Analysis | Peak latency (ms) | Onset latency of first cluster (ms) | Offset latency of last cluster (ms) | Correlation (Spearman's R) |
|---|---|---|---|---|
| **A. Category representations shared between infants and adults** | | | | |
| Infants | 200 (200–360) | 160 (-200–160) | 540 (540–920) | 0.61 (0.49–0.72) |
| Adults | 120 (120–1,000) | 100 (60–100) | 1,000 (900–1,000) | 95.67 (92.46–97.84) |
| Difference | 80 (-800–200) | 60 (-300–80) | n/a | n/a |
| **B. Category representations shared between infants and adults (based on the decoding accuracy, averaged adult RDMs)** | | | | |
| Infants | 340 (-180–980) | 280 (-140–520) | 340 (280–1,000) | |
| Adults | 420 (-180–660) | 400 (-60–680) | 580 (560–1,000) | |
| Difference | -80 (-480–1,000) | -120 (-460–360) | n/a | |
| **C. Category representations shared between infants and adults (based on the correlation, 1 – Pearson's *R*)** | | | | |
| Infants | -20 (-140–340) | 180 (-180–180) | 660 (660–1,000) | |
| Adults | 840 (140–960) | 80 (40–100) | 1,000 (1,000–1,000) | |
| Difference | -860 (-1,000–140) | 100 (-260–120) | n/a | |
| **D. Category representations shared between infants and adults (based on the Euclidean distance)** | | | | |
| Infants | 320 (-200–40) | 160 (-200–160) | 440 (440–780) | |
| Adults | 140 (120–980) | 100 (60–120) | 1,000 (880–1,000) | |
| Difference | 180 (-680–180) | 60 (-320–80) | n/a | |

**Table S2. Statistical Details of the Across-Population Representational Similarity Analysis, related to Figure 2 and Figure S2.** The table enumerates the conducted analysis in the 1$^{st}$ column (from **A** to **D**), the peak latency within a significant cluster (2$^{nd}$ column), the onset latency of the first significant cluster (3$^{rd}$ column), the offset of the last significant cluster (4$^{th}$ column), and correlation at peak (5$^{th}$ column). Notes: All values are averages across subjects (infant $n$ = 40, adult $n$ = 20), with 95% confidence intervals are reported in brackets.

| Comparison | | Infants | | Adults | |
|---|---|---|---|---|---|
| | | Correlation (Spearman's *R*) | Significance (*P* –value) | Correlation (Spearman's *R*) | Significance (*P* –value) |
| **A. Correlations between neural (based on 95% CIs definition) and model RDMs and significance** | | | | | |
| **1) At the whole-model level** | | | | | |
| Gabor filter | | 0.1764* | <0.001 | 0.5252* | <0.001 |
| VGG | | 0.1250* | 0.0029 | 0.4898* | <0.001 |
| AlexNet | | 0.1207* | 0.0228 | 0.4583* | <0.001 |
| ResNet | | 0.1221* | 0.0292 | 0.4640* | <0.001 |
| **2) At the layer-group-level in deep neural networks (DNNs)** | | | | | |
| VGG layer group | 1 | -0.0121 | 0.8833 | -0.0096 | 0.5190 |
| | 2 | 0.0993 | 0.0373 | 0.4184* | <0.001 |
| | 3 | 0.1493* | 0.0068 | 0.5156* | <0.001 |
| | 4 | 0.1493* | 0.0068 | 0.5156* | <0.001 |
| | 5 | 0.1207 | 0.0449 | 0.4583* | <0.001 |
| | 6 | -0.1450 | 0.0704 | -0.1271 | 0.0731 |
| | 7 | 0.0436 | 0.6149 | 0.3492* | 0.0022 |
| AlexNet layer group | 1 | -0.0993 | 0.0950 | -0.2692* | 0.0051 |
| | 2 | -0.0050 | 0.9721 | 0.2577* | 0.0227 |
| | 3 | 0.0736 | 0.1363 | 0.4123* | <0.001 |
| | 4 | 0.1493* | 0.0068 | 0.5156* | <0.001 |
| | 5 | 0.0679 | 0.3208 | 0.3750* | 0.0013 |
| | 6 | 0.0679 | 0.3208 | 0.3750* | 0.0013 |
| | 7 | -0.0121 | 0.7799 | 0.2402* | 0.0022 |
| ResNet layer group | 1 | 0.1964* | 0.0163 | 0.3538* | 0.0075 |
| | 2 | 0.1979* | 0.0066 | 0.4833* | <0.001 |
| | 3 | 0.1464 | 0.0473 | 0.4479* | <0.001 |
| | 4 | 0.0707 | 0.3279 | 0.3865* | <0.001 |
| | 5 | 0.0707 | 0.3279 | 0.3865* | <0.001 |
| **B. Correlations between neural (based on cluster-extent definition of shared representations) and model RDMs and significance** | | | | | |
| **1) At the whole-model level** | | | | | |
| Gabor filter | | 0.1222* | 0.0210 | 0.6686 | <0.001 |
| VGG | | 0.1073* | 0.0171 | 0.6514 | <0.001 |
| AlexNet | | 0.1036 | 0.0745 | 0.5686 | <0.001 |
| ResNet | | 0.0951 | 0.0443 | 0.6200 | <0.001 |
| **2) At the layer-group-level in deep neural networks (DNNs)** | | | | | |
| VGG layer group | 1 | -0.0301 | 0.5909 | -0.0257 | 0.3985 |
| | 2 | 0.0973 | 0.1251 | 0.5943* | <0.001 |
| | 3 | 0.1151* | 0.0101 | 0.6743* | <0.001 |
| | 4 | 0.1151* | 0.0101 | 0.6743* | <0.001 |

| | | | | | |
|---|---|---|---|---|---|
| | 5 | 0.1036 | 0.0745 | 0.5686* | <0.001 |
| | 6 | -0.0315 | 0.8107 | -0.2114* | 0.0024 |
| | 7 | 0.0764 | 0.3379 | 0.4171* | <0.001 |
| AlexNet layer group | 1 | -0.0796 | 0.3023 | -0.4371* | <0.001 |
| | 2 | 0.0502 | 0.6163 | 0.3886* | <0.001 |
| | 3 | 0.0794 | 0.1861 | 0.5743* | <0.001 |
| | 4 | 0.1151 | 0.0101 | 0.6743* | <0.001 |
| | 5 | 0.0842 | 0.2362 | 0.4400* | <0.001 |
| | 6 | 0.0842 | 0.2362 | 0.4400* | <0.001 |
| | 7 | 0.0449 | 0.6737 | 0.2571* | 0.0042 |
| ResNet layer group | 1 | 0.0771 | 0.3800 | 0.4457* | <0.001 |
| | 2 | 0.1050 | 0.2041 | 0.6000* | <0.001 |
| | 3 | 0.0900 | 0.1086 | 0.5829* | <0.001 |
| | 4 | 0.0673 | 0.2169 | 0.5429* | <0.001 |
| | 5 | 0.0673 | 0.2169 | 0.5429* | <0.001 |

**Table S3. Correlations between Neural and Model RDMs and Significance, related to Figure 3 and Figure S3.** The table reports the correlation between model and neural RDMs (defined by temporal extent of 95% CIs around the peak) and computational model RDMs at the **(A)** whole model and **(B)** layer-group level and its significance (infant $n$ = 40, adult $n$ = 20, two-tailed sign permutation tests, $P$ < .05, FDR-corrected, significant correlations are indicated by a star).

| Analysis | Latency (ms) | | | Frequency (Hz) | | | Peak decoding accuracy (%) or correlation (Spearman's R) |
|---|---|---|---|---|---|---|---|
| | Peak | Cluster onset | Cluster offset | Peak | Cluster onset | Cluster offset | |
| **A. Spectral characterization of infant category representations** | | | | | | | |
| Infants | 400<br>(160–580) | 180<br>(-80–340) | 520<br>(440–960) | 4.63<br>(2.91–6.73) | 3.85<br>(2.65–4.63) | 6.13<br>(4.22–8.91) | 55.21 (52.38–57.62) |
| Adults | 180<br>(180–200) | -100<br>(-100– -100) | 1,000<br>(1,000–1,000) | 5.59<br>(5.09–6.13) | 2.00<br>(2.00–2.00) | 30.00<br>(30.00–30.00) | 95.66 (93.79–97.28) |
| **B. Similarity between infant theta-based category representations and adult category representations resolved in time and frequency (based on cluster-extent definition)** | | | | | | | |
| | 120<br>(120–380) | 80<br>(-20–120) | 180<br>(180–1,000) | 17.13<br>(8.12–20.65) | 12.95<br>(2.00–14.21) | 22.67<br>(20.65–30.00) | 0.67 (0.54–0.79) |
| **C. Similarity between infant theta-based category representations and adult category representations resolved in time and frequency (based on 95% confidence interval definition)** | | | | | | | |
| | 120<br>(120–380) | 80<br>(-20–100) | 180<br>(180–1,000) | 17.13<br>(9.78–20.65) | 12.95<br>(2.00–14.21) | 22.67<br>(20.65–30.00) | |
| **D. Variability in spectral power peak frequency and peak latency in the infant and adult sample** | | | | | | | |

| | Peak latency | | | | Peak frequency | | |
|---|---|---|---|---|---|---|---|
| | Range (ms) | Variance | Interquartile range (ms)** | | Range (Hz) | Variance | Interquartile range (Hz)** |
| Infants | 0–980 | 7122.8 | 150 | | 3.19–30 | 30.16 | 1.38 |
| Adults | 100–560 | 898.95 | 20 | | 6.13–8.12 | 0.41 | 0.96 |

**Table S4. Statistical Details of Spectral Analyses and the Variability in Spectral Power, related to Figure 4 and Figure S4.** The table enumerates the statistical details for spectral analysis and the variability in spectral power for infants and adults. The conducted analysis was shown in 1st column. For **(A–C)**, we listed the peak latency within a significant cluster (2nd column), the onset latency of the first significant cluster (3rd column), and the offset of the last significant cluster (4th column), the peak frequency within a significant cluster (5th column), the onset frequency of the first significant cluster (6th column), the offset frequency of the last significant cluster (7th column), and decoding accuracy or correlation at peak (8th column), 95% confidence intervals are reported in brackets. All values are averages across subjects (infant n = 40, adult n = 20), with 95% confidence intervals in brackets. For **(D)**, we listed the range, variance and interquartile range of the peak frequency and peak latency in the spectral power for both infants and adults. Consistently across all the three listed measures, for both time and frequency dimensions, adults show lower variability than infants. **Interquartile range is the difference between the 75th and 25th percentiles.
